# Dengue virus 2 lacking N153 glycosylation displayed enhanced recognition by neutralizing antibodies

**DOI:** 10.64898/2026.07.31.741530

**Authors:** Guntur Fibriansah, Thiam-Seng Ng, Xin-Ni Lim, Aaron W.K. Tan, Donald Heng Rong Ting, Gavin R. Screaton, James E. Crowe, Sylvie Alonso, Shee-Mei Lok

## Abstract

Deglycosylated (N153Q) dengue virus (DENV) mutant shows attenuated infection in a mouse model, mainly due to its increased antibody susceptibility. This is consistent with the neutralization assay showing human monoclonal antibodies 2D22 and C10 are more potent towards the mutant than the wild-type (WT) virus. Here, we compared the cryoEM structures of WT and mutant viruses complexed with these Fabs (2.7-3.2 Å resolution). We observed increased occupancies for both Fabs on the mutant virus, suggesting higher accessibility of epitopes that were previously blocked by glycosylation. Using biolayer interferometry, we showed although the Fabs have slower binding rate to the mutant than WT virus, they also have slower dissociation rate. The slow dissociation rate might contribute to higher Fab occupancies, as once bound, they remain associated with the virus. N153Q mutant might be a good vaccine candidate, as important epitopes are made more accessible for stimulating highly potent antibodies.

## Introduction

Dengue virus (DENV), which consists of four serotypes (DENV1-4), is a positive-sense RNA virus. It belongs to the *flaviviridae* family and other important related pathogens are yellow fever, tick-borne encephalitis, West Nile, and Zika viruses. Dengue is spread primarily by *Aedes aegypti* mosquito. DENV is endemic in the tropical and subtropical regions^1,2^. The number of DENV infection cases is estimated to be nearly 390 million annually, including approximately 100 million cases of dengue fever and 500,000 cases of dengue haemorrhagic fever or dengue shock syndrome, resulting in 500,000 hospitalizations and 20,000 deaths^3^.

There are no highly effective vaccines for dengue, and vector control is challenging. A safe and effective dengue vaccine that stimulates balanced immune responses against all four serotypes would be ideal to stop disease transmission. Two dengue vaccines have been licensed: Dengvaxia®, and Qdenga, developed by Sanofi and Takeda Pharmaceutical Company, respectively. Dengvaxia demonstrated particularly lower efficacy for the DENV2 serotype^4,5^ and has shown to cause serious adverse effects in vaccinees without prior exposure to any dengue serotypes, limiting its use to seropositive individuals^6^. Qdenga has shown higher and more balanced vaccine efficacy than Dengvaxia^7,8^. However, serotype-specific negative vaccine efficacy was observed in the vaccinated seronegative individuals^9^, suggesting that long-term surveillance of Qdenga vaccine recipients is required^10^.

The DENV particle contains the nucleocapsid - plus-sense RNA genome wrapped together with capsid proteins. The nucleocapsid is then contained within a bilayer lipid membrane, and facing the exterior and anchored on the membrane are the E and M proteins^11-^_13_. The E protein is the main antigenic protein stimulating antibody response^14^. The E ectodomain consists of three domains-domain I (DI), DII, and DIII. This is followed by trans-membrane helices anchoring the E protein to the lipid bilayer membrane^13^. The E proteins on virus surface forms homodimers (**Fig. S1**). Three E protein homodimers are stacked parallel together forming a raft structure, and the raft structures are organized into a herringbone pattern on the virus surface^11–13^. There are in total 180 copies of E proteins (**Fig. S1**) arranged in an icosahedral symmetry. Each asymmetric unit (ASU) contains three E proteins-molecules (mols) A, B, and C, located near 5-, 2-, and 3-fold vertices, respectively (**Fig. S1**). A raft structure contains two adjacent ASUs (**Fig. S1**).

Unlike other orthoflaviviruses, which have only one glycosylation site at residue N153/N154, DENV has two highly conserved N-glycosylation sites on the E protein, at residues N67 and N153 (**Fig. S1**)^15–17^. Glycosylation on the E protein likely plays significant roles in the viral infection, transmission and pathogenesis^18^. N67 glycosylation has been shown to be the binding site for Dendritic Cell-Specific Intercellular adhesion molecule-3-Grabbing Non-integrin (DC-SIGN) ^19,20^. DENV lacking glycosylation at N67 or N153, or both sites maintained its ability to replicate and propagate in mosquito cells^20^. However, the ablation of N67 glycosylation impaired the assembly of new infectious particles in mammalian cells, whereas the removal of N153 glycosylation resulted in a virus with reduced infectivity^20,21^. Although the virus lacking N67 glycosylation (N67Q mutation) could propagate in mosquito cells, another mutation (K64N) was observed near to the original glycosylation site, highlighting the importance of glycosylation at this site^21^. The removal of glycosylation at residue 153 has been shown to increase the membrane fusion pH threshold^22,23^.

In other viruses, such as SARS-CoV-2, human immunodeficiency virus and influenza virus, the glycans on the surface proteins can be used to shield important or functional epitopes from antibodies^24–26^. Ting *et al.* reported that the removal of DENV2 N153 glycosylation site enhanced antibody-mediated viral clearance^27^. Consistent with what others have found^20^, DENV2 lacking the N153 glycosylation site (N153Q mutant) exhibited only mildly impaired infectivity in Vero and Huh-7 cell lines^27^. However, in a symptomatic mouse model of severe dengue, the N153Q mutant virus was significantly attenuated, N153Q virus experienced accelerated IgM-mediated clearance^27^.

In this study, we used two highly potent human monoclonal antibodies (HMAb) (2D22 and C10) ^28,29^ whose epitopes have been shown to be in the vicinity of the N153 glycosylation site, to study by cryoEM their binding characteristics to DENV2 N153Q mutant compared to fully glycosylated DENV2 WT virus. We showed that both HMAbs have different binding modes towards DENV2 WT and N153Q mutant viruses. HMAb 2D22 is a DENV2-specific antibody, whereas HMAb C10 is a cross-neutralizing antibody that neutralizes DENV1-4 as well as ZIKV^28–30^. The two antibodies were shown to bind to E-protein dimer epitopes that consist of DII of one E protein and DIII or DII/DI-DII hinge region/DIII, respectively, of the other E protein within a dimer^29,30^. Here, we have improved the resolution of the cryoEM maps of the Fab complexed with WT and mutant viruses to ∼2.8A compared to previous structures (2D22, 6 Å and C10: ∼3.5 Å resolutions), thus showing atomic interactions. Our *in vitro* neutralization tests confirmed that IgG 2D22 and C10 are more potent towards the N153Q mutant than the WT virus as reported by Ting *et al.*^27^. This is consistent with our cryoEM structures showing higher Fab occupancies for both 2D22 and C10 on the N153Q mutant. We also performed biolayer interferometry to further understand the binding kinetics of the two antibodies to DENV2 WT and N153Q mutant. Overall, it suggests that in the N153Q mutant virus, the epitopes recognized by highly potent antibodies are much more accessible, thus making it a better vaccine candidate for stimulating highly potent antibodies.

## Results

### DENV2 N153Q mutant is structurally stable at 37°C

Some DENV2 strains (NGC and 16681) can change from smooth to bumpy surfaced particles when incubation temperature is raised from 4°C to 37°C^31,32^. Here we examined whether temperature changes induce structural changes to DENV2 (D2Y98P) and its deglycosylated counterpart N153Q mutant. CryoEM images of the DENV2 WT and the N153Q mutant at 4°C showed that most of the particles had a round, smooth surface morphology, indicating mature dengue particles (**Fig. S2A**). Only a few spiky-looking particles, which is indicative of the immature or partially immature particles, were observed. Incubation of viruses at 37°C for 30 min did not change the morphology (**Fig. S2A**), suggesting both WT and N153Q mutation are structurally stable. The cryoEM maps of DENV2 WT and the N153Q mutant have been determined to a resolution of 3.2 Å and 2.7 Å, respectively (**Figs. 1A and S3A, and Table S1**). At these resolutions, the cryoEM maps show most of the amino acid residue side chains (**Fig. 1B**). Superposition of the WT and N153Q mutant structures shows that the two structures are identical (**Fig. 1C**), thus indicating that N153 glycans do not influence the formation and stability of the overall structures of the virus surface.

**Figure 1.**
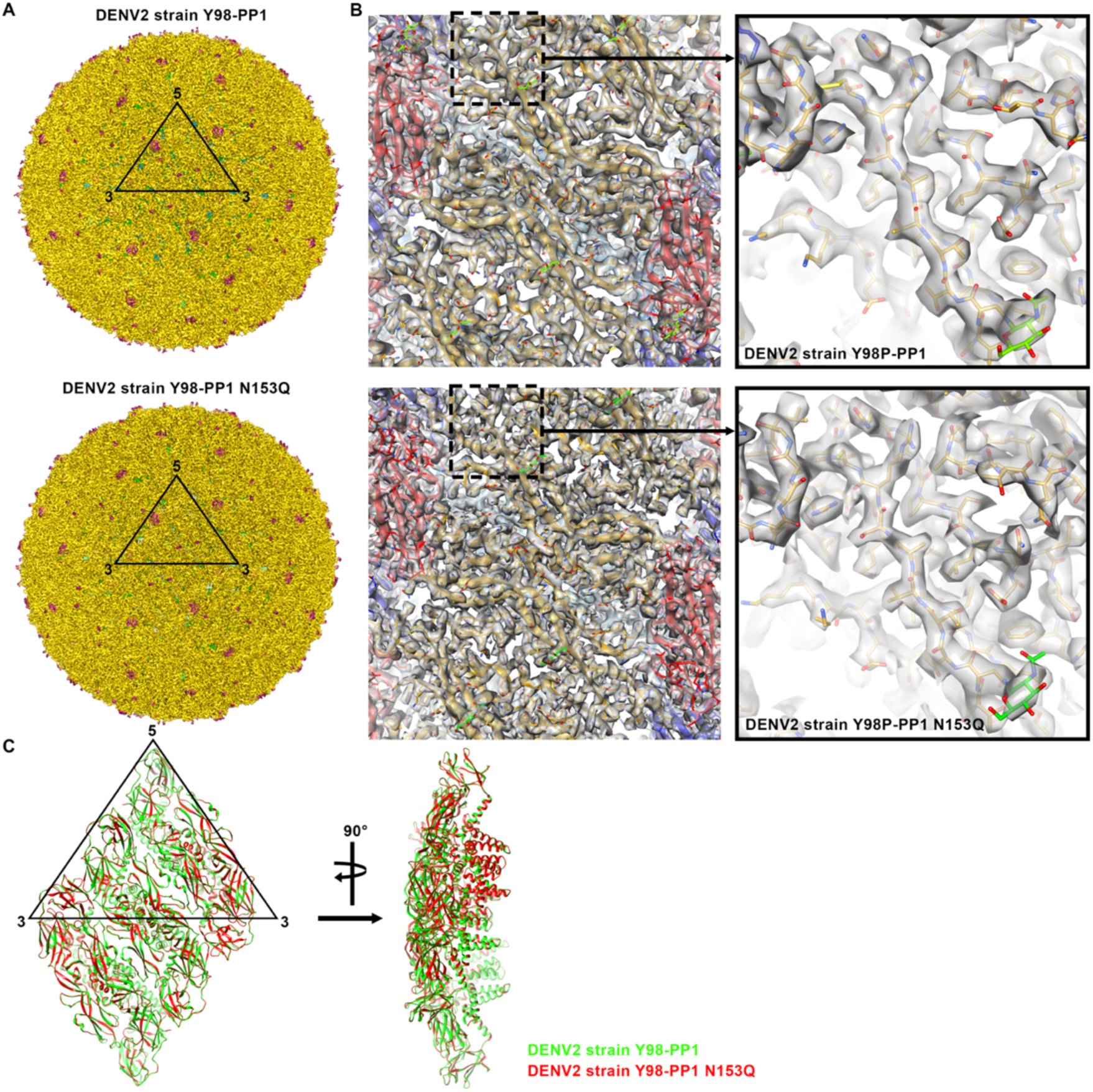
The cryoEM structures of DENV2 WT and N153Q mutant at 4°C. (**A**) The cryoEM maps of the DENV2 WT and N153Q mutant were determined to resolutions of 3.2 and 2.8 Å, respectively. The maps were color-coded radially: green for 150-200 Å, yellow for 201-234 Å, and magenta for > 235 Å. In the maps, the ectodomain part of E proteins appears in yellow, the lipid bilayer membrane (visible between E protein densities) appears in green, and the protruding part of DIII of E protein molecule B and some glycosylation, particularly on N67, appear in magenta. The cryoEM maps of DENV2 WT and N153Q mutant show the same pattern, suggesting they are structurally identical. An icosahedral ASU is indicated by a black triangle. (**B**) The high-resolution cryoEM maps display side chain densities, allowing for more accurate model coordinate fitting of the amino acid residues. The E protein structure is shown in ribbon representation, with DI, DII, and DIII colored in red, yellow, and blue, respectively. The cryoEM densities are displayed as transparent grey surfaces. (**C**) Superposition of the structures of the DENV2 WT and the N153Q mutant shows that the two structures are identical.

### The glycan loop of the DENV2 N153Q exhibited higher mobility

The glycosylation at residue N153 of DENV2 WT is visible from the cryoEM densities on the residues (**Fig. S4A**). The glycan loops (residues 140-160) of the three E proteins (mols. A, B, and C) in an icosahedral ASU have well-defined densities, with the amino acid side-chains visible. However, in contrast, the corresponding glycan loops in the N153Q mutant have poorer densities, particularly around residue H158 (**Fig. S4B**). The cryoEM map confirms the N153Q mutation, as there is an absence of glycan densities. The poorer density of the glycan loops in the N153Q mutant suggests that the glycan residues at N153 play a part in stabilizing the conformation of the glycan loop. Further analysis of the B-factor distribution shows that the glycan loops of the three E proteins in the N153Q mutant structures have higher B-factors compared to those of the wild type (**Fig. S4**). The B-factor distribution of the rest of the E protein structures in both structures is similar, they both show overall low B-factors on the E protein ectodomain and high B-factors on the tip of the transmembrane helices. This suggests a higher mobility of the glycan loop in the N153Q mutant structure.

### DENV2 N153Q mutant is more pH sensitive

A virus aggregation assay was conducted to assess the pH at which the virus starts to expose the fusion loop, as occurs during the fusion process. The exposure of the fusion loop results in the aggregation of the virus due to hydrophobic interactions between the fusion loops and viral membranes. DENV2 WT started to aggregate at pH 6.0, while DENV2 N153Q aggregated at the higher pH of 6.5 (**Fig. 2A**). Consistent with previous literature^22,23^, the increased pH threshold in this aggregation assay indicates that DENV2 N153Q might undergo endosome membrane fusion at an elevated pH.

**Figure 2.**
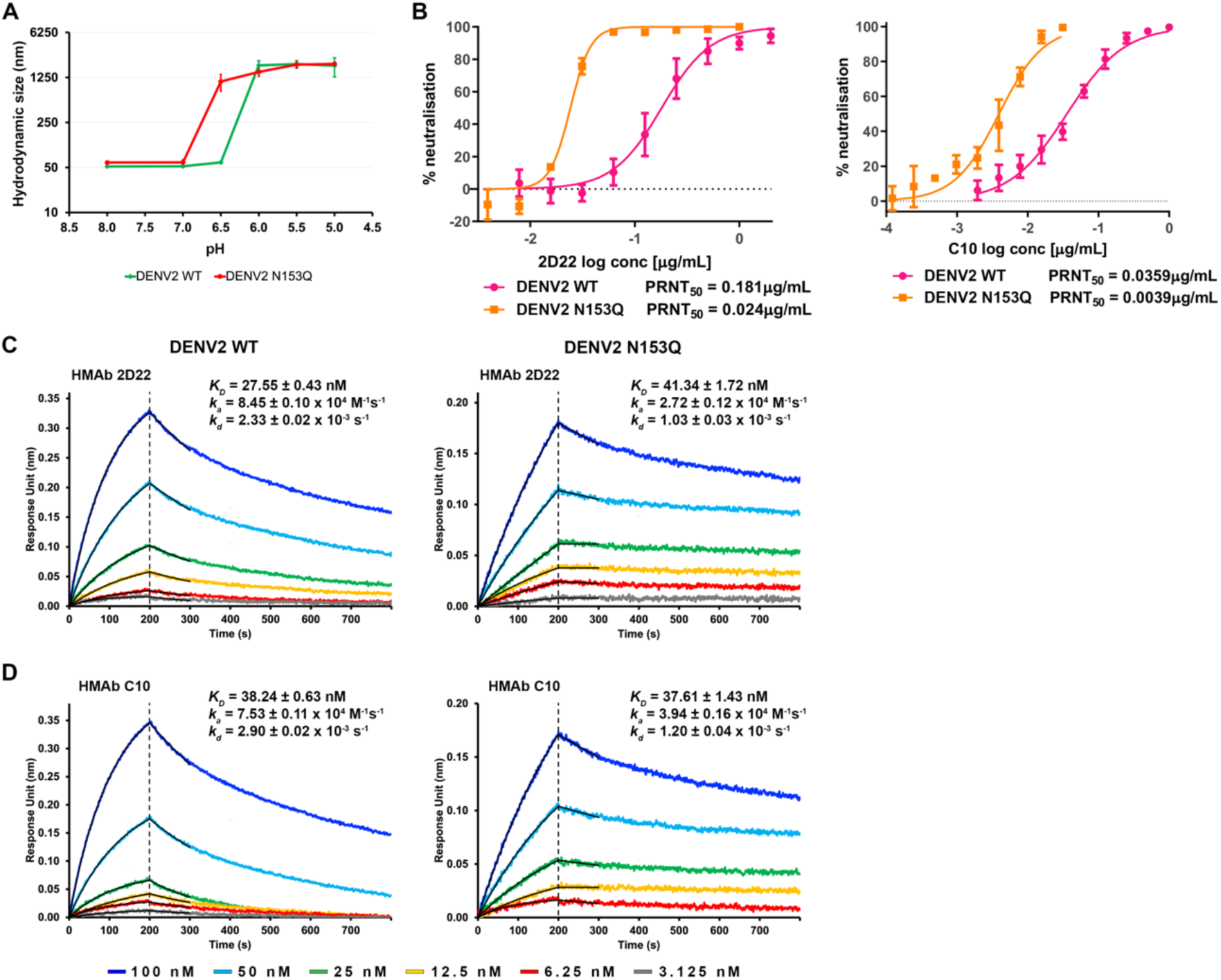
DENV2 N153Q aggregates at higher pH than WT and HMAbs 2D22 and C10 neutralized the DENV2 N153Q more effectively. (**A**) DENV2 N153Q started to aggregate at a higher pH threshold than DENV2 WT. The virus aggregation is due to the exposure of the E protein fusion loop, hence these results indicated that DENV2 N153Q likely undergoes fusion at higher pH threshold that DENV2 WT. (**B**) The plaque reduction neutralization test conducted on DENV2 WT and N153Q mutant using HMAbs 2D22 (left) and C10 (right) revealed that the concentration of antibody required to neutralize 50% (PRNT_50_) of the N153Q mutant virus was lower than that of the wild type for both HMAbs 2D22 and C10. Error bars represent standard deviation. (**C**-**D**) HMAbs 2D22 and C10 bound to DENV2 with similar K_D_ values. Removing the glycan at N153 resulted in slower binding of the antibody to the virus (lower in k_a_, association rate value); however, it also reduced the rate at which the antibody dissociates from the virus (lower in k_d_, dissociation rate value). The combination of both actions resulted in minimal changes in K_D_ values. This suggests that the presence of glycan at N153 in the WT virus assists in opening the epitope, thus accelerating the antibody binding process. In the N153Q mutant, once the antibody is bound, it remains tightly associated with the virus. The binding affinity was measured by biolayer interferometry method.

### HMAbs 2D22 and C10 neutralize the N153Q mutant more efficiently than the wild type

Consistent with data generated by Ting *et al.*^27^, plaque reduction neutralization test on DENV2 WT and N153Q mutant with HMAbs 2D22 and C10 showed that the antibodies neutralized N153Q mutant better than WT (**Fig. 2B**). HMAb 2D22 neutralized the N153Q mutant with a PRNT_50_ of 0.024 μg/mL, which is 7.5 times better than to the WT virus (PRNT_50_ = 0.181 μg/mL). Similarly, HMAb C10 neutralized the N153Q mutant 9.2 times more efficiently than WT DENV2 (PRNT_50_ of 0.0039 μg/mL vs 0.0359 μg/mL, respectively). The lower PRNT_50_ values of DENV2 N153Q were not due to the attenuation of the virus, since N153Q mutant virus was shown to retain parental *in-vitro* fitness in BHK-21 cells – the same cells used for the PRNT assay here - as reported by Ting *et al.* ^27^.

### HMAbs 2D22 and C10 have reduced association and dissociation rates towards DENV2 N153Q

The decrease in the PRNT_50_ values of HMAbs 2D22 and C10 against DENV2 N153Q mutant could be due to an increase in the binding affinity of antibodies to E protein, leading to an increase in Fab occupancy on the virus surface. To investigate this, we performed a biolayer interferometry assay to measure the affinity of HMAbs 2D22 and C10 to DENV2 WT and the N153Q mutant. The measurement results showed that HMAb 2D22 bound to DENV2 N153Q with an association rate 3.1x poorer than DENV2 WT (**Fig. 2C**). A similar trend was also observed with HMAb C10, showing a 1.9x poorer association rate compared to WT virus (**Fig. 2D**). However, once the antibodies have bound to the N153Q particle, they remain more tightly bound when compared to WT virus. The dissociation rate of HMAbs 2D22 and C10 in the binding experiment with DENV2 N153Q is 2.3x and 2.4x slower, respectively, than that with DENV2 WT (**Fig. 2C-D**). Decreases in both dissociation and association rates resulted in a moderate 1.5 times poorer (higher K_D_ value) affinity of HMAb 2D22 to DENV2 N153Q compared to its affinity to DENV2 WT, while the binding affinities of HMAb C10 to DENV2 WT and N153Q mutant are similar.

### More efficient binding of Fabs 2D22 and C10 to the N153Q mutant virus

HMAb 2D22 has been previously shown to bind to DENV2 strains PVP94/07 and NGC. The structure of DENV2 strain PVP94/07 complexed with Fab 2D22 at 4°C was previously determined to a resolution of 6.5 Å^29^, whereas that of DENV2 NGC complexed with Fab C10 at 28°C to 3.6 Å resolution^30^. Strain PVP94/07 has a very tight E-E protein interactions on the virus surface whereas, that of the NGC virus strain are much looser and hence when heated to 37°C, NGC strain shows the bumpy surface structure^29,31^.

Here we examine the interactions of Fabs 2D22 and C10 binding to another DENV2 WT strain (D2Y98P) and its N153Q mutant at 4 and 37 °C (**Fig. S2B-C**). To investigate how the N153Q mutation increases the neutralization efficiency of HMAbs 2D22 and C10, we have determined the high resolution cryoEM structures of the WT and the N153Q mutant in complex with Fab 2D22 or C10 at 37°C, which is the physiological human body temperature (**Figs. 3 and S3B-C, and Table S2**). The structures of DENV2 WT and N153Q mutant in complex with Fab 2D22 were determined to 2.8 Å resolution, while those in complex with Fab C10 to 2.9 Å resolution (**Figs. S3B-C and Table S2**). The cryoEM maps of the DENV2 WT and N153Q in complex with Fab 2D22 showed that the Fab occupancies in both complex structures appeared to be the same (**Fig. 3**). However, this differs for the Fab C10, where the WT virus complex showed lower Fab occupancies - 120 Fabs compared to the 180 on the N153Q mutant (**Fig. 3**).

**Figure 3.**
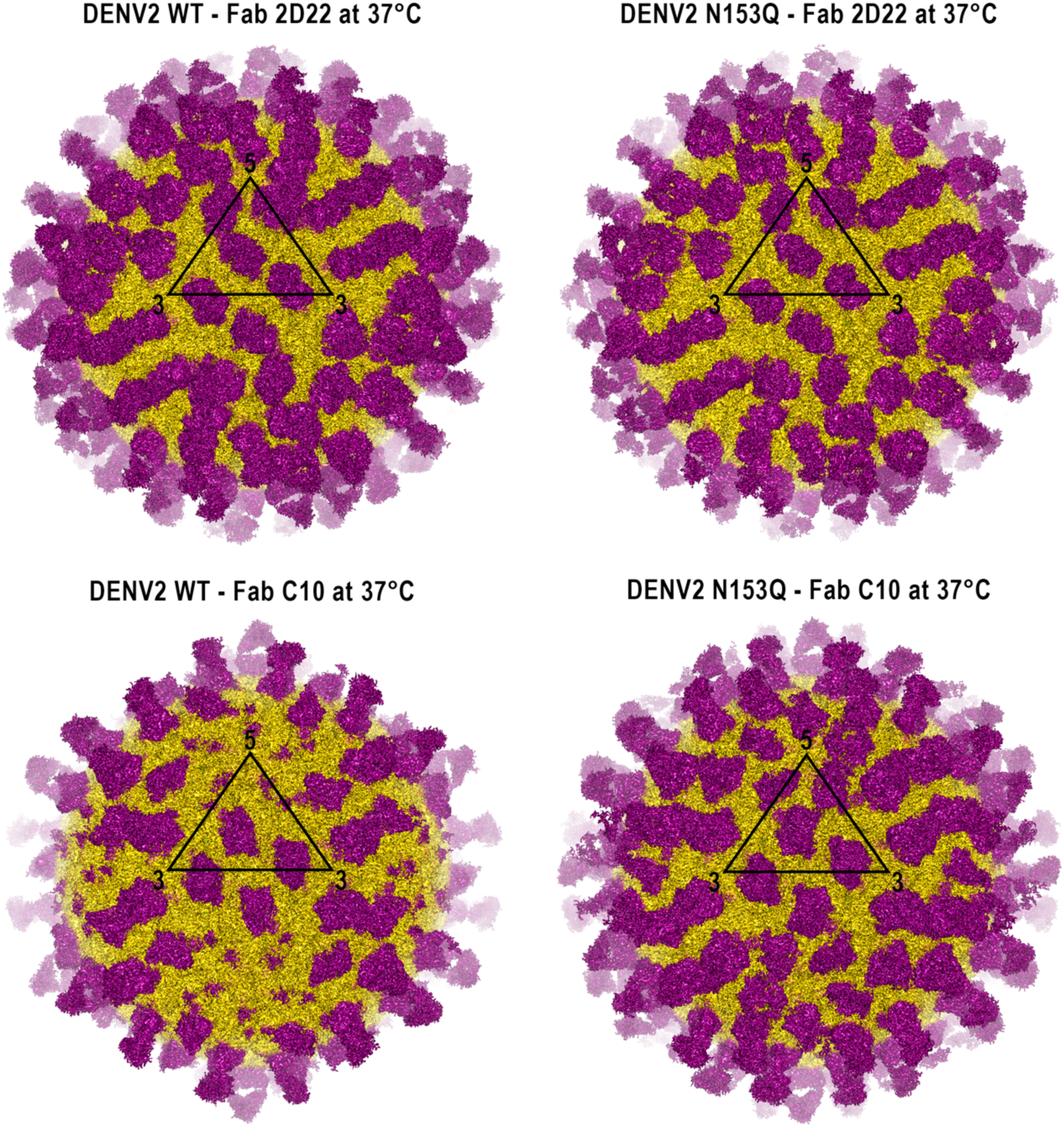
More efficient binding of Fab 2D22 and C10 to DENV2 N153Q mutant. CryoEM maps of the DENV2 WT and N153Q mutant complexed with Fab 2D22 or C10. The cryoEM maps were color-coded as described in Fig. 1A. The densities of Fabs 2D22 and C10 are shown as magenta surfaces. The black triangle indicates an icosahedral ASU. There are 180 copies (3 Fab/ASU) of Fabs 2D22 bound to both the wild type and N153Q mutant virus particles. Fab C10, on the other hand, shows higher occupancy on the N153Q mutant than WT virus – 180 copies versus 120 (2 Fabs/ASU) copies of Fabs.

The 2D22 and C10 epitopes derived from our high resolution maps, are roughly consistent with the previously published lower resolution maps^29,30^. In general, 2D22 epitope comprised of DII of one E protein and the DIII of the other E protein within an E protein dimer, along with a few additional residues from the adjacent E protein dimer^29^. Whereas, the epitope of HMAb C10 was located on the DII of one E protein and the DII, DI-DII hinge region, and DIII of the other E protein within the E protein dimer, also with a few additional residues from the adjacent E protein dimer^33^.

Comparison of the Fabs binding epitopes between our high resolution DENV2 WT and the N153Q mutant maps showed largely similar interacting residues for each Fabs (**Fig. 4A-B, Table S3-S13**). However, we also observed that both Fab 2D22 and C10 have slightly more interactions with the E proteins on the mutant virus surface. The number of interactions engaged by Fabs 2D22 and C10, on the E protein dimer and also to adjacent E protein increases by 2-4 interactions in the DENV2 N153Q mutant compared to DENV2 WT, except for Fab 2D22 bound near the 3-fold vertices (**Table S14**). We observed from the cryoEM maps of DENV2 WT and N153Q mutant in complex with Fab 2D22 or C10, while the Fab densities are strong, the nearby E protein glycan loops densities are much poorer (**Fig. S5A**). This suggests that there are no specific interactions between them. It might also suggests that the glycan loops has been pushed aside and have adopted different conformations and hence being averaged out during the image reconstruction process. The E protein molecule A in the WT - Fab C10 complex did not have Fab C10 bound to this epitope and hence, the glycan loop densities were strong (**Fig. S5A**).

**Figure 4.**
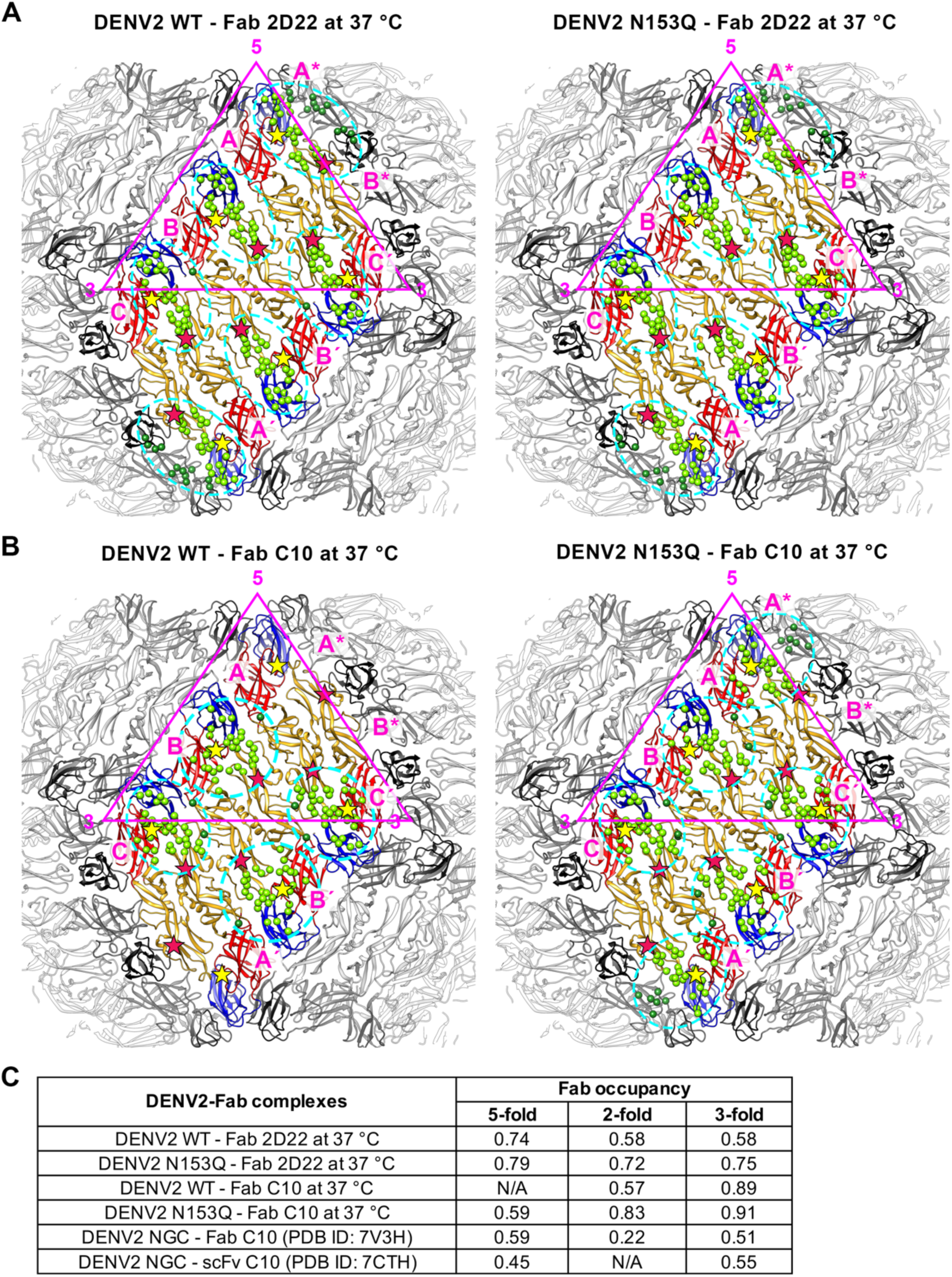
Footprints of Fabs 2D22 and C10 on the E proteins on DENV2 N153Q mutant and WT virus are largely the same, except occupancies for both Fabs are consistently higher on the N153Q mutant. (**A-B**) The 2D22 and C10 epitopes on DENV2 WT and N153Q mutant were identified using the program “CONTACT” from the CCP4 Program Suite ^53^ set with a threshold distance of 4.2 Å. HMAbs 2D22 and C10 epitopes are circled with cyan dashed lines. The epitope residues on an E protein dimer and the adjacent dimer are represented by light and dark green spheres, respectively. The glycosylation at residues N67 and N153 is indicated by red and yellow stars, respectively. DI, DII, and DIII colored in red, yellow, and blue, respectively, in one E protein raft, while the others rafts are colored in shades of of grey. An icosahedral ASU is indicated by a dark pink triangle. (**C**) The table shows occupancies of Fabs 2D22 and C10 on the three individual epitopes in an ASU. Fab occupancy is estimated by calculating the ratio between the average map values at the paratope residues and the values at the epitope residues [(Fab ÷ E protein dimer) x 100 %]. The occupancy of Fab C10 or scFv C10 on another DENV2 strain (NGC) was taken from the previous publications ^30,39^.

To further investigate the effect of the N153Q mutation on Fab binding, we calculated the occupancy of Fabs 2D22 and C10 on each epitope by comparing the Fab densities to the E protein densities. The results showed that the Fab occupancy on the N153Q mutant is consistently higher than on the WT virus (**Fig. 4C**), suggesting that the removal of glycosylation site at N153Q improves Fab binding. Specifically, Fab 2D22 occupancy at epitopes near the 2- and 3-fold vertices increased significantly from 0.58 for both epitopes in DENV2 WT to 0.72 and 0.75, respectively, in DENV2 N153Q mutant. This occupancy level is similar to that of Fab 2D22 at epitopes near the 5-fold vertices, where it was 0.74 in DENV2 WT and only slightly increased to 0.79 in DENV2 N153Q mutant. The Fab C10 occupancy at epitopes near the 2- and 5-fold vertices showed a significant increase from 0.58 and 0 (no binding), respectively, in DENV2 WT to 0.83 and 0.59, respectively, in DENV2 N153Q mutant. The Fab C10 occupancy at epitope near the 3-fold vertices is the highest (0.89) compared to the other epitopes in DENV2 WT, with only a minimal increase to 0.91 in the DENV2 N153Q mutant.

The superposition of the structures of Fab 2D22 or C10 bound to DENV2 WT and the respective complexes with the N153Q mutant showed that both structures, except for the glycan loop parts, are essentially identical. This suggests that the N153Q mutation did not alter the binding conformation of both antibodies (**Fig. S5B**). To investigate the differences in the binding mechanisms of Fabs 2D22 and C10 on DENV2 WT and the N153Q mutant, leading to differing Fab occupancies, the DENV2 WT and N153Q E protein dimer-Fab complexes for each Fab near the 2-, 3-, and 5-fold vertices were aligned by superimposing the distal end of DII (residues 65-119 and 239-251), a region containing epitope residues. To describe the relative Fab binding orientation with respect to the E protein dimer, the binding axis for each bound Fab was measured. The binding axis is defined as the vector that connects the center of the epitope of HMAb 2D22 or C10 (i.e., Cα atom of residue W101 or K246, respectively) to the center of the core β-sandwiches of the HMAb 2D22 or C10 V^H^ and V^L^ domains (i.e. the center between Cα atoms of L45 of the heavy chain and K46 of the light chain, or Q39 of the heavy chain and Q40 of the light chain, respectively).

The binding axes of Fabs 2D22 and C10, when bound to their respective epitopes on the E proteins of DENV2 WT and N153Q, are shown in **Fig. 5**. There are two groups of Fab 2D22 binding axes, the first group consists of Fab bound to epitopes near 2- and 3-fold vertices of DENV2 WT and N153Q mutant, and the second group consists of Fab bound to epitopes near the 5-fold vertices (**Fig. 5A**). Since the majority of the epitopes residues for Fab 2D22 are located on the DII of one E protein in a dimer (**Fig. 4A, and Tables S3-S5, S8-S10**), the presence of these two distinct groups of binding axes indicates there are two structurally different epitope on E protein dimer. These two epitopes differ based on the relative position of DII of one E protein and the DIII of the other E protein and/or adjacent E protein. Even though there are differences in the local structures of the epitopes, Fab 2D22 was still able to bind all epitope suggesting the flexibility of Fab paratope in recognizing the epitope residues. Notably, the distance between axes of the three Fab 2D22 binding sites on DENV2 N153Q is closer than those on the DENV2 WT indicating the N153Q reduced the local structure differences between the three epitopes on the virus (**Fig. 5A**). This resulted in similar Fab 2D22 occupancy at the three epitopes near to 2-, 3-, and 5-fold vertices in the DENV2 N153Q mutant, with occupancy level of 0.79, 0.72, and 0.75, respectively.

**Figure 5.**
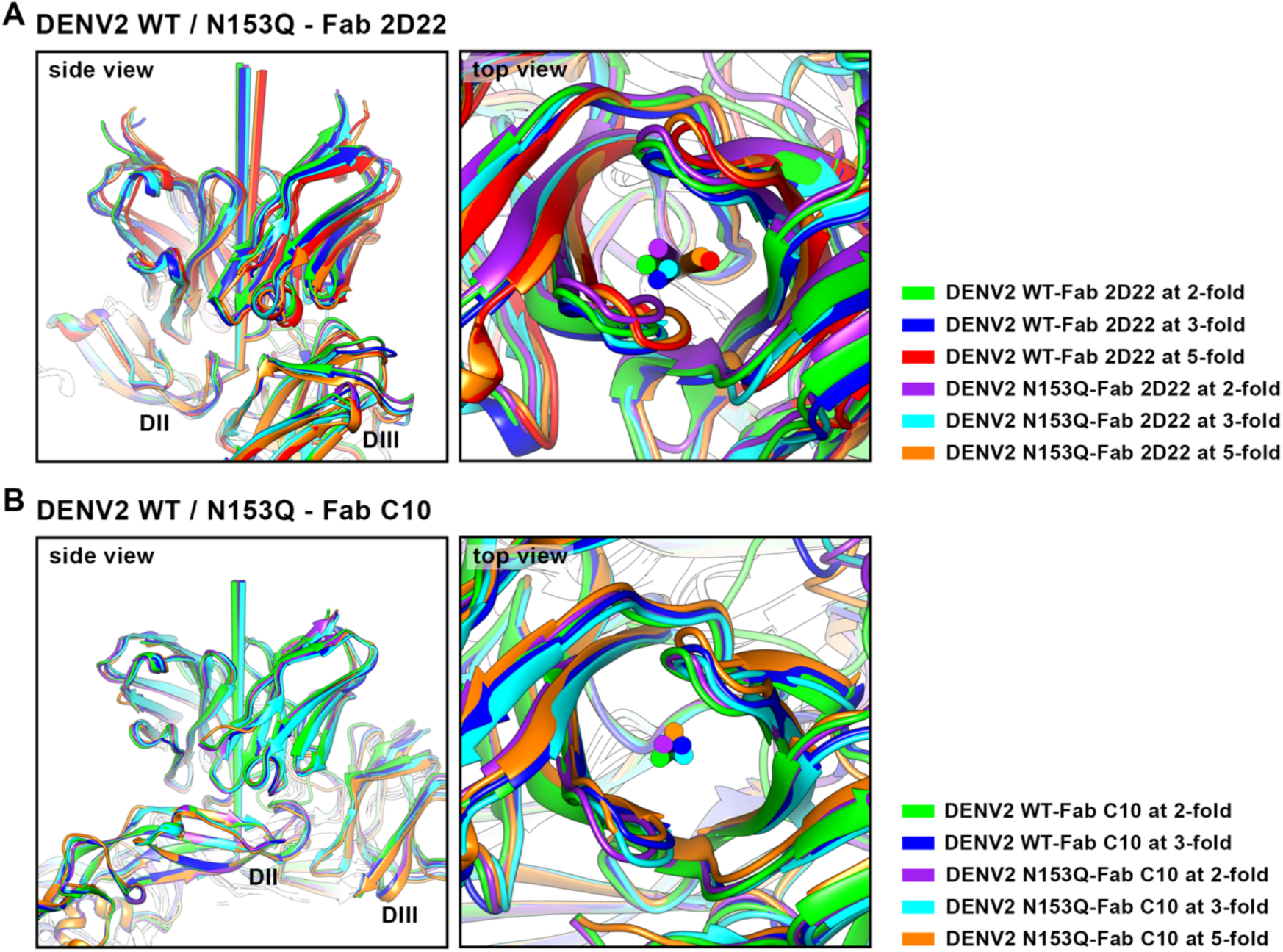
HMAb 2D22 has more flexible binding axes compared to those of HMAb C10. (**A**) The binding axes of Fab 2D22 on DENV2 WT and N153Q are distributed into two groups: one group consists of Fabs bound near the 5-fold vertices, and the other groups consists of those bound near the 2-fold and 3-fold vertices. The difference in binding axes becomes less pronounced in the Fabs bound to DENV2 N153Q, as indicated by the smaller angle difference between the axes. (**B**) The binding axes of Fab C10 on DENV2 WT and N153Q are distributed as one tight group with minimal differences in binding axes. The binding axis is defined as the vector that connects the center of the epitope of HMAb 2D22 or C10 (i.e., Cα atom of residue W101 or K246, respectively) to the center of the core β-sandwiches of the HMAb 2D22 or C10 V^H^ and V^L^ domains (i.e. the center between Cα atoms of L45 of the heavy chain and K46 of the light chain, or Q39 of the heavy chain and Q40 of the light chain, respectively). The DENV2 WT and N153Q E protein dimer-Fab complexes were aligned by superimposing the distal end of DII (residues 65-119 and 239-251), which contains epitope residues.

Different from Fab 2D22, Fab C10 binds to the epitopes near the 2-, 3-, and 5-fold vertices of DENV2 WT and the N153Q mutant in a more unified manner. The interacting residues in the epitope of Fab C10 are distributed almost equally on DII of one E protein and DII/DI-DII hinge region/DIII of the other E protein (**Fig. 4B, Tables S6-S7, S11-S13**). The Fab C10 binding axes are clustered into a single group, indicating a stricter structural requirement of the epitope for Fab binding (**Fig. 5B**). The epitope located near the 5-fold vertices of DENV2 WT likely does not fit the structural requirement, hence there is no Fab C10 binding at this epitope. However, this epitope on DENV2 N153Q mutant was able to adopt a conformation that permitted Fab C10 binding. Superposition of the DII of E protein structures of uncomplexed and Fab C10-complexed DENV2 WT and N153Q near the 5-fold vertices showed that the DIII of the E protein in DENV2 N153Q mutant has a different conformation, moving away from the Fab C10 bound position (**Fig. S6B**). This suggests that DENV2 N153Q exhibits higher structural flexibility compared to DENV2 WT, allowing such movement and accommodating the Fab C10 binding at this epitope.

## Discussion

The N153Q mutation on E protein from DENV2 strain D2Y98P did not affect virus structure thermal stability. The mutant virus maintained its smooth surface morphology after 30 min incubation at 37°C. Exposure of N153Q mutant to different low pH buffers showed the virus particles have aggregated at higher pH conditions than the WT virus. The low pH induced E proteins to flip up and expose their fusion loops for interaction with membranes of adjacent virus particles thereby causing virus-virus aggregation. This mimics the fusion of virus with endosomal membrane event. Results therefore suggest the DENV2 N153Q mutant undergoes virus fusion at a higher pH threshold than that used by the WT virus, hence fusion might happen in an earlier endosome compartment. The histidine residue is likely critical for pH sensing and the initiation of fusion due to its pKa of approximately 6.0, which is the pH where fusion occurs^34^. Several histidine residues are identified as playing an important role in the fusion process: (1) H317 at the DI/DIII interface^35–37^, (2) H144, at the DI/DIII interface^36^, (3) H282 at the DI/stem helices interface^37^. The change in the protonation state of these residues at lower pH alters the E protein conformation, leading to the exposure of the fusion loop. The residue H144 is located in the glycan loop, which shows increased mobility in the N153Q mutant. Changes in local conformational and the microenvironment, such as water accessibility, can influence the properties of affected residues^38^. Hence, the increased mobility of the glycan loop in DENV2 N153Q mutant might affect the pKa of H144 possibly resulting in the protonation of this residue at a higher pH, leading to the exposure of the fusion loop and membrane fusion compared to that of DENV2 WT.

The presence of glycans at N153 on another DENV2 strain (PVP94/07) in complex with Fab 2D22 was observed to cause conformational changes in the glycan loop^29^. This glycosylation likely causes steric hindrance for Fab binding to the epitope^29^. The binding of Fab 2D22 to our DENV2 WT (D2Y98P) also caused a shift in the glycans and hence also the glycan loop (**Fig. S5A**). However, in the N153Q mutant virus, which lacks glycosylation at this position, structural changes were still observed in its glycan loops for both Fabs 2D22 and C10 complexed virus structures (**Fig. S5A**). This observation suggests that the glycan loop itself likely also partially blocks access to the epitope. Superposition of their E proteins of the uncomplexed DENV2 WT and N153Q mutant onto the respective E proteins in the Fab 2D22 or C10 complexed virus structures shows that the glycan loop residues, with or without glycosylation at residue 153, will clash with the Fab molecules (**Fig. 6A-B**). Thus, the glycan loop changes its conformation upon the binding of these antibodies onto the E proteins, regardless of the glycosylation at residue 153 (**Fig. S5A**).

**Figure 6.**
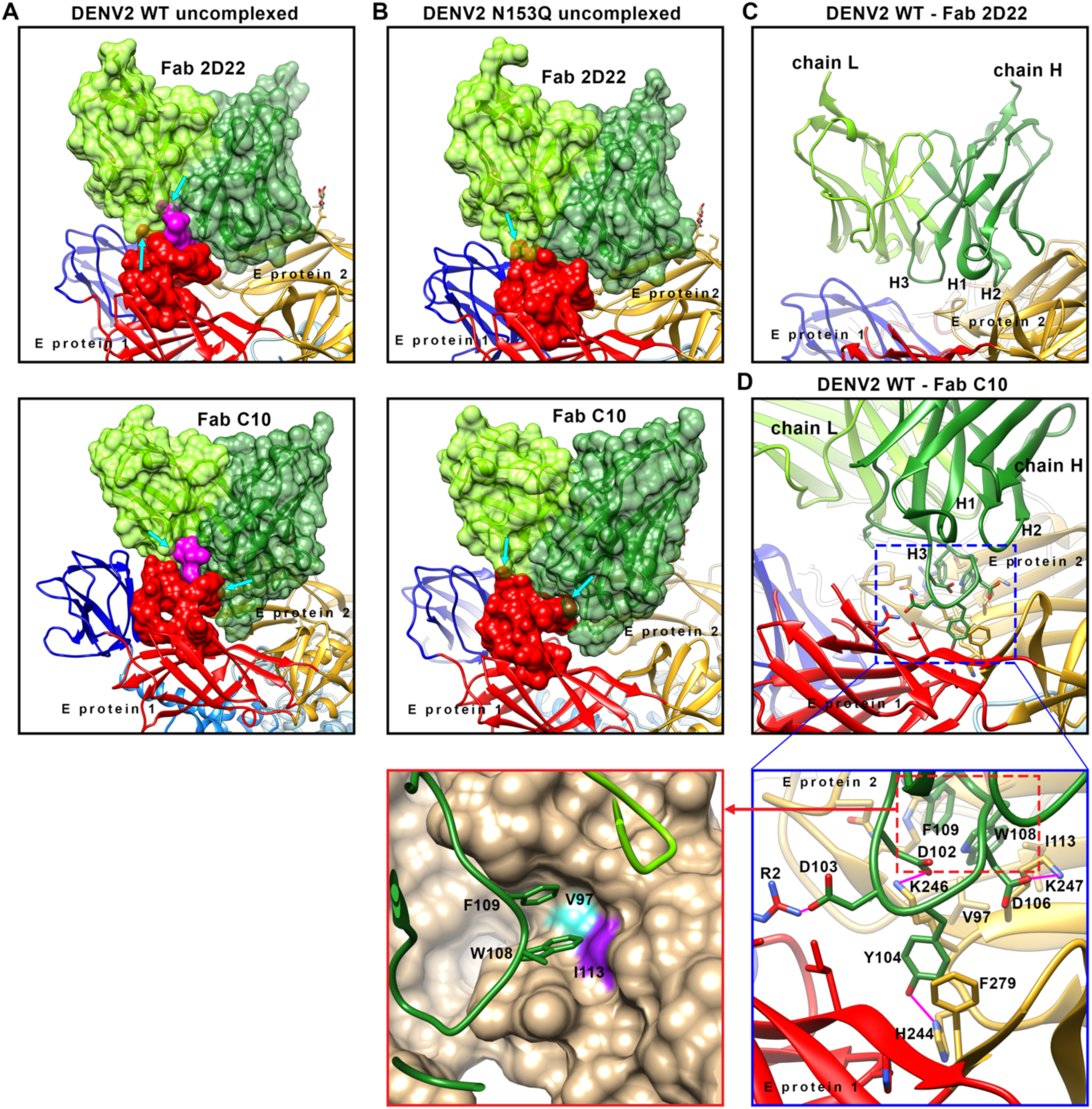
Superposition of Fabs 2D22 and C10 complex structures onto uncomplexed WT shows clashes with the glycan loop and glycan residues on the uncomplexed WT. The E proteins of the uncomplexed DENV2 WT and N153Q mutant were superimposed onto the respective E proteins of the Fab 2D22 or C10 complex structures (**A**-**B**). (**A**) Both the glycan loop and glycan residues attached to N153 in the uncomplexed DENV2 WT clash with Fabs 2D22 and C10. This suggests Fabs 2D22 and C10 likely induced conformational changes to the glycan loop upon binding. The clashes are indicated by cyan arrows. (**B**) Likewise, the glycan loop in the uncomplexed DENV2 N153Q mutant clashes with the bound Fab 2D22 or C10, leading to movement of this loop in the complexed structure. The E proteins of the Fab 2D22 or C10 complexed virus structures are not shown, whereas the E proteins of the uncomplexed virus structures are shown in ribbon representation, with DI, DII, and DIII colored in red, yellow, and blue, respectively. The glycan loop (residues 140-160) is shown in solid surface representation, while the heavy and light chains of the Fab molecules are shown in ribbon representation with a transparent surface, colored dark and light green, respectively. The glycan residues are shown in magenta solid surface representation. CryoEM structures of the WT and N153Q mutant viruses complexed with (**C**) Fab 2D22 and (**D**) Fab C10 near the 3-fold vertices are shown. The CDR-H3 loop of Fab C10 is longer compared to that of Fab 2D22. (**D**) (**Top**) Upon binding to DENV2, the CDR-H3 loop of Fab C10 is inserted between DI of one E protein monomer and DII of another E protein monomer. (**Bottom, right**) Zoom-in view of the interactions between the H3 loop residues and the epitope residues. (**Bottom, left**) Fab residues W108 and F109 interact with E protein residues V97 and I113 in a DII pocket, respectively. The CDR-H1, -H2, and -H3 of Fabs 2D22 and C10 are indicated. The E protein dimer in the bottom-left panel of (**D**) is shown as a light-brown surface, with residues V97 and I113 colored in cyan and purple, respectively.

The biolayer interferometry analysis showed that HMAbs 2D22 and C10 bind to the DENV2 N153Q mutant with a lower association rate compared to their binding to DENV2 WT. This might suggest the bulkiness of the glycan in WT virus could help move the glycan loop out of the way thereby allowing easy access of antibodies to 2D22 and C10 epitopes. Whereas, the antibodies might have a harder time avoiding the flexible glycan loop in the N153Q mutant, leading to a lower association rate. However, once bound, the Fabs bind more tightly to N153Q mutant, as indicated by a lower dissociation rate. This is corroborated by a general increase in the number of interactions made between the Fab and the E protein dimer (**Table S14**).

Our cryoEM structure of the Fabs complexed with WT virus shows minimal interactions with the glycan. The BLI data shows both antibodies have better association rate to WT virus particle but they also dissociate faster than the mutant. Using all atom MD simulation, Ting *et al*.^27^ reported that the glycans at both N67 and N153 had transient interactions with residues within 2D22 epitope. We speculate that these interactions are weak and likely do not represent a major obstacle for the high affinity antibody to displace the glycan interactions with the amino acids within the 2D22 epitope. We instead propose that the bulkiness of the glycan help to get the glycan loop out of the way for Fabs to bind, which may thus explain the better association rate constant for Fabs to WT virus than the N153Q mutant virus. Of note, the biolayer interferometry experiment was performed using virus that was grown in insect C6/36 cells; therefore, it has less bulky glycans attached to the N-glycosylation site compared to that grown in mammalian cells. The affinity parameters, including the association rate constant, for the binding of antibodies to virus grown in mammalian cells such as Vero and Huh-7 might be different. The neutralization activity of antibodies to DENV2 WT grown in different cell lines has been demonstrated to be lower in virus with bulkier glycans grown in mammalian cells^27^, indicating weaker antibody binding affinity.

The MD simulations of the E protein dimer complexed with Fab 2D22 showed that Fab likely bound to DENV2 WT poorly (higher RMSD) than that to the N153Q mutant across all cell lines^27^. When bound to WT virus grown in mammalian cell lines (Vero and Huh-7), MD simulations showed Fab bound less tightly to the E protein dimer compared to WT virus from insect C6/36 cells^27^. However, the simulation was performed on an E protein dimer already bound to Fabs, hence it simulating the dissociation process rather than association. Thus, it indicates that the bulkier the glycans, the faster Fab 2D22 would dissociate, therefore it aligns well with our BLI data well. Additionally, MD simulations studied the interactions between E protein dimer in soluble environment with antibodies, which may not fully recapitulate the interactions occurring in the context of the whole virus particle. Consistently, we observed differences in Fab occupancies to different dimers within a raft on the virus likely due to differences in the curvature and local contacts^30^. We also observed some interactions of Fab across dimers. Our high resolution maps showed more residues on the N153Q mutant interacting with the Fabs than the WT virus, this might explain for the slower dissociation rate.

Although the binding affinities of HMAbs 2D22 and C10 to the DENV2 WT and DENV2 N153Q mutant are similar, neutralization assays showed that the antibodies neutralize the DENV2 N153Q mutant more effectively than DENV2 WT. It is possible that the lack of glycans leads to increased exposure of some partially hidden epitopes. Therefore, compared to WT virus, we observed an overall increase in Fab occupancies for both Fabs (**Fig. 3**) in our cryoEM maps. We also observed different levels of exposure of the epitopes in the three different E protein molecules within an icosahedral ASU when comparing the occupancy between different Fabs (**Fig. 4C).** This suggests that even though the binding affinities are similar, the exposure of each epitope, which leads to stronger interactions between Fab and E proteins, plays a more important role in neutralization efficiency than the rate at which the Fab binds to the epitope.

The Fab 2D22 occupancies across the three E proteins are similar, particularly on DENV2 N153Q mutant, whereas for Fab C10, the highest and lowest Fab occupancies are on the epitopes near the 3-fold and the 5-fold vertices, respectively. This suggests that there is another factor that determines Fab occupancies besides enhanced epitope accessibility. It has been demonstrated that Fab C10 can cause conformational changes leading to overall increased particle dynamics and distortion in DENV2 strain NGC^30^. The cryoEM structure of DENV2 strain NGC in complex with Fab C10 showed that the Fab C10 occupancy is lower for the epitope located near the 2-fold vertices. However, for our DENV2 WT, the lowest occupancy is observed for the epitope near the 5-fold vertices. This suggests that DENV2 exhibits a different particle dynamic motion upon binding of Fab C10 compared to DENV2 strain NGC.

The Fab C10 binding axes on E protein of DENV2 WT or N153Q has been shown to have a similar binding orientation (**Fig. 5B**). Superposition of the E proteins with Fab bound near the 2-, and 3-fold vertices in the structures of Fab C10-complexed DENV2 WT, DENV2 N153Q, and DENV2 NGC (PDB ID 7V3H), as well as scFv (single chain variable fragment) C10 complexed DENV2 NGC-2 (a substrain of DENV2 NGC with some mutations, **Fig. S7**) (PDB ID: 7CTH), showed that the Fab C10 binding axes at these two epitopes are closely clustered into one group (**Fig. S6A**). However, the binding axes of Fab C10 bound to the epitope near the 5-fold vertices are divergent.

Fab C10 bound to the epitope near the 2-fold vertices of DENV2 strain NGC with limited occupancy of 0.22 (**Fig. 4C**) ^30^, whereas scFv C10 was unable to bind to this position on DENV2 NGC-2^39^. When the E proteins of Fab C10-complexed DENV2 NGC, scFv C10-compexed DENV2 NGC-2, and uncomplexed DENV2 WT and N153Q were superimposed at the epitope near the 2-fold vertices (**Fig. S6B,** top right panel), the DIII of DENV2 NGC-2, in which scFv C10 does not bind to, deviates from the other structures (indicated by red arrows). The E protein near the 2-fold vertices of DENV2 WT and N153Q did not undergo conformational changes when Fab C10 bound this epitope (**Fig. S6B,** top left panel) with occupancy of 0.57 and 0.83, respectively. This indicate that the structural conformation of the epitope on DENV2 WT is suitable for Fab binding and N153Q mutation enhanced the Fab binding. In contrast, the epitope on DENV2 NGC-2 requires conformational changes to accommodate Fab C10 binding. It is possible that E protein near the 2-fold vertices in DENV2 NGC originally had the same structure as the DENV2 NGC-2 but then changed its conformation for Fab C10 binding. DENV2 NGC-2 is not as flexible due to some mutations on the E protein compared to DENV2 NGC (**Fig. S7**). The E proteins of DENV2 NGC at this location have been shown to be more dynamic and led to an overall virus surface morphological changes at 37°C^31,40^. Other DENV2 NGC substrains that have mutations, such as DENV2 NGC-2, have been shown to be less dynamic and did not change morphology at higher temperature^41^.

Comparison of the E proteins near the 5-fold vertices of Fab C10 complexed DENV2 NGC and scFv C10 complexed DENV2 NGC-2 with those of uncomplexed DENV2 WT and N153Q mutant also indicates conformational changes in the DIII upon the binding of Fab C10 or scFv C10 (**Fig. S6B**, middle right panel, blue arrows). This is similar to the changes observed in the Fab C10 binding to DENV2 N153Q mutant (**Fig. S6B**, middle left panel, black arrows).

Fab 2D22 bound to the three epitopes on DENV2 WT or N153Q mutant with minimal conformational changes (**Fig. S8**), whereas Fab C10 binding to epitope near the 5-fold vertices in DENV2 N153Q mutant or near the 2-fold vertices in DENV2 NGC requires some structural changes (**Fig. S6B**), suggesting a unique binding mechanism displayed by Fab C10. The complementarity-determining region of an antibody consists of H1-H3 loops in the heavy chain and L1-L3 loops in the light chain. A comparison of these loops in Fabs 2D22 and C10 showed that the H3 loop (residues 101-113) of Fab C10 is longer than that of Fab 2D22 (**Fig. 6C-D**).

In the complex with the E protein dimer, the H3 loop is inserted between DI of one E protein and DII of the other E protein (**Fig. 6D**). The residues of the H3 loop make numerous interactions, including salt bridges between D102, D103, and D106 of Fab C10 and K246 (DII - E protein monomer 2), R2 (DI - E protein monomer 1), and K247 (DII - E protein monomer 2), respectively (**Fig. 6D**, bottom right panel**, Tables S6-S7**). Additionally, residue Y104 of Fab C10 forms a hydrogen bond and hydrophobic interaction with H244 (DII - E protein monomer 2) and F279 (DI - E protein monomer 1), respectively (**Fig. 6D**, bottom right panel). Notably, residues W108 and F109 of Fab C10 interact with V97 and I113 (DII - E protein monomer 2), which are slightly hidden on the E protein dimer surface, forming a pocket (**Fig. 6D**, bottom left panel). Residue R2 is located under the N153 glycan loop and is exposed only when the glycan loop is removed upon Fab C10 binding. The specific interactions of the H3 loop of Fab C10 with these residues on DI and DII indicate the important roles of this loop in recognizing the binding site. The binding of Fab C10 might require conformational changes to accommodate the H3 loop; otherwise, this loop could become a hindrance to binding.

The lack of glycans at N153Q has been shown to increase epitope exposure, hence increasing the occupancy of Fabs 2D22 and C10. These antibodies, known as EDE antibodies, are highly neutralizing^28,29,42^. The N153Q mutation does not alter the virus stability, yet it allows increased exposure of epitopes combined with increased dynamics of the E proteins on the viral surface leading to enhanced binding of EDE antibodies. This could explain the increased susceptibility of the N153Q mutant to IgG-mediated neutralisation observed by us and *Ting et.al.*^27^. Therefore, using DENV2 N153Q as an attenuated virus vaccine may help induce a higher immune response specifically targeting the EDE antibody epitope.

## Materials and Methods

### Virus sample preparation

The previously described protocol ^12,43^ was followed to grow and purify the viruses. *Aedes albopictus* C6/36 cells were used to propagate the viruses, and these cells were grown in RPMI medium supplemented with 10% FCS until they reached approximately 80% confluency. Following that, the cells were inoculated with DENV2 WT (D2Y98P) (GenBank accession number: JF327392) or N153Q mutant^27^ at a multiplicity of infection of 0.1 and incubated at 28°C for 4 days. To eliminate cell debris, the virus in the supernatant was collected through centrifugation at 9,000 x g for 30 min. Subsequently, the virus presents in the supernatant underwent precipitation by adding 8% w/v polyethylene glycol 8000 in NTE buffer (10 mM Trs-HCl pH 8.0, 120 mM NaCl, and 1 mM EDTA), followed by an overnight incubation at 4°C. The resulting precipitated virus was then collected through centrifugation at 14,300 x g. After that, the virus pellet was resuspended in NTE buffer and further purified by subjecting it to centrifugation through a 24% w/v sucrose cushion. The virus was then layered on top of a linear gradient of potassium tartrate ranging from 10% to 30% w/v. After extracting the band that contained the virus, it was buffer exchanged into NTE buffer and concentrated using an Amicon Ultra-4 centrifugal concentrator (Millipore). The centrifugal concentrator had a 100 kDa molecular weight cut-off membrane. All purification procedures were performed at a temperature of 4°C. The purified virus was then stored at this temperature until it was frozen on cryoEM grids. To determine the purity and concentration of the purified virus, SDS-PAGE with Coomassie staining was used. The concentration of the E protein was estimated by comparing the intensity of the band to a range of different concentrations of bovine serum albumin protein standard.

### Virus aggregation assay

One microliter of purified DENV2 WT and N153Q mutant with an E protein concentration of 0.25 mg/mL was diluted in 19 μL assay buffer that consists of 150 mM NaCl and 20 mM MES pH 5.0, 5.5, 6.0, 6.5, sodium phosphate pH 7.0, or Tris pH 8.0. The sizes of the virus at different pH levels were measured by the dynamic light scattering method using Zetasizer Nano S machine (Malvern) at 25°C. The sample was pre-incubated inside the cuvette chamber for 5 min prior to each measurement. The results were analyzed using Zetasizer Nano software version 8.02. The experiment was done in triplicate.

### Plaque reduction neutralization test (PRNT)

The neutralization activities of the HMAb 2D22 and C10 on both DENV2 WT and N153Q mutant were determined by PRNT. Two-fold serially diluted HMAb 2D22 samples with starting concentration of 4 µg/mL and 2.0 µg/mL were added to equal and fixed quantity of DENV2 WT and N153Q viruses, respectively, and then the mixtures were incubated at 37°C for 30 min. Whereas for HMAb C10, the starting concentrations used were 2.0 µg/mL and 62.5 ng/mL, respectively, and the experiments were performed the same as for HMAb 2D22. One hundred microlitres of each mixtures were then layered on BHK-21 cells in a 24-well plate and incubated at 37°C for 1 h. The infected cells were subsequently washed, overlaid with carboxyl-methyl cellulose, and incubated at 37°C. Cells were fixed and stained with crystal violet after 4 days, and the number of plaques forming units (PFU) were counted. Percentage neutralization was determined from the comparison of the reduction in the number of PFU in specific antibody dilutions to the respective positive control (without antibody). PRNT_50_ is the concentration of the antibody wherein there is a 50% reduction of PFU. This was determined using nonlinear regression in GraphPad Prism 9. The experiments were performed three times, each with triplicate samples. The mean values are shown as data points and the standard deviations are indicated as error bars.

### Biolayer interferometry

The binding affinities of HMAbs 2D22 and C10 to DENV2 WT and N153Q were measured by biolayer interferometry using an Octet RED96e (Sartorius) with anti-human IgG Fc capture biosensors (AHC biosensors, Sartorius). The tips of the AHC biosensors were first hydrated in a Tris buffer containing 20 mM Tris (pH 8.0), 150 mM NaCl, and 0.025% Tween-20 for 1 h at room temperature. The HMAbs 2D22 and C10, along with DENV2 WT and N153Q, were also diluted in this Tris buffer. The DENV2 WT and N153Q were two-fold serially diluted, with final concentrations of the E protein ranging from 0.78 to 100 nM (0.17 to 5.42 μg/mL), while antibody IgGs were diluted to 10 μg/mL.

Two hundred microliters of solutions (DENV2 WT or N153Q, HMAb 2D22 or C10 IgG, and buffers) were transferred into 96-well flat bottom black plates (Corning). The plate was incubated at a temperature of 30°C and shaken at 1000 rpm throughout the experiment. The experiments began with a baseline measurement of each biosensor tip in the Tris buffer. The tips were subsequently placed into wells containing the antibodies for antibody immobilization until achieving a response of approximately 0.5-0.6 nm. After antibody immobilization, the tips were dipped in the Tris buffer for 1 min to remove excess IgG antibodies. This was followed by a 200 s baseline measurement in other wells containing Tris buffer.

The association step was performed on the serially diluted DENV2 WT or N153Q for 200 s, followed by dipping the tips back into the Tris buffer used in the previous baseline measurement for a 600 s dissociation step. After dissociation, the biosensor tips were regenerated in 10 mM Glycine (pH 2.0) to remove bound IgG. Baseline subtraction was conducted with tips dipped into the Tris buffer without DENV2 WT or N153Q. A 1:1 kinetic model fitting was used to calculate the binding affinity of each antibody and virus combination. The experiment was performed at least three times for each combination of antibody and virus, and a representative sensorgram is shown.

### Fabs 2D22 and C10 preparation for cryoEM studies

HMAbs 2D22 and C10 IgG were digested with immobilized papain protease (Thermo Fisher Scientific) according to the manufacturer’s instructions. Briefly, the antibodies were digested using papain with a weight ratio of papain:IgG of 1:100 and incubated at 37°C overnight. The papain beads were then removed from the supernatant using centrifugal filter unit (Millipore), and the supernatant was buffer exchanged to a 20 mM Tris buffer at pH 8.0. The Fab was purified using a 1 mL Resource Q anion exchange column (Cytiva). The Fab 2D22 and C10 were obtained in the flow-through fractions. The fractions containing the Fab were buffer exchanged to a 20 mM Tris-HCl buffer at pH 8.0 and 150 mM NaCl, and then concentrated using an Amicon Ultra-4 centrifugal concentrator (Millipore) with a 10 kDa molecular weight cut-off membrane.

### Cryo-EM sample preparation

The purified DENV2 WT and N153Q viruses were mixed with Fabs 2D22 or C10 at a molar ratio of E protein to Fab of 1:1.5 and incubated at 4°C for 30 min. The virus-Fab complex samples were then further incubated at 37°C for an additional 30 min, followed by another incubation at 4°C for approximately 2 h before being frozen. Uncomplexed viruses at 37°C were prepared in the same manner, but without the addition of Fab. For sample preparation, a 2.1 μL sample was applied to either a glow-discharged Quantifoil R 2/1 plus C2 on 300 copper mesh or ultrathin carbon film on a lacey carbon support film 400 mesh copper grid (Ted Pella). The sample was applied using a Vitrobot Mk IV (Thermo Fisher Scientific) operated at 4°C and 100% humidity. After application, the sample was blotted with filter paper and immediately plunged into liquid ethane. It was then kept at liquid nitrogen temperature until it was ready to be imaged.

### CryoEM imaging

Single-particle cryoEM image acquisition was performed using SerialEM software ^44^ on a Titan Krios cryogenic transmission electron microscope (Thermo Fisher Scientific). The microscope was equipped with a field emission gun operating at 300 kV, a K3 direct electron detector, and a Gatan imaging filter (GIF) post-column energy filter (Gatan). The micrographs were acquired in counting mode at a nominal magnification of 64,000, corresponding to 1.345 Å per pixel on the camera. Beam image shift in SerialEM was utilized to target holes in a 3-by-3 multishot pattern after each stage movement, except for the DENV2 WT at 4°C images, which were collected using a single-shot method. The number of micrographs, dose rate, total electron exposure, and defocus range can be found in **Tables S1** and **S2**.

### Cryo-EM image processing

The micrographs of the DENV2 wild type at 4°C, which belong to a single optics group, were processed directly in RELION ^45^. The remaining micrographs were sorted into nine optics groups based on beam image-shift values. Movie frames were aligned in RELION, and the contrast transfer function (CTF) parameters were estimated using CTFFIND4 ^46^. Manual particle picking was performed on a subset of datasets, and the particles were extracted and binned by a factor of 4 for initial 2D classification. Good 2D classes were selected and used as templates for automated particle picking in RELION. The automatically picked particles were further extracted and binned by a factor of 4 for 2D classification. After manually selecting the good 2D classes, the particles were subjected to 3D classification with particle alignment. Icosahedral symmetry was imposed during the 3D classification, and the cryoEM map of DENV4 (EMDB 2485) was used as the reference map. Particles from the good 3D classes were selected and subjected to 3D refinement. Further refinements were performed using particles extracted from unbinned images. The 3D image reconstruction was improved by conducting CTF refinement and Bayesian polishing in RELION. The final cryoEM maps were then subjected to B-factor sharpening in RELION. The resulting cryoEM maps are shown in **Figs. 1 and 3**. The resolution of each cryoEM map was estimated using the Fourier shell correlation (FSC) curve with a cutoff value of 0.143. The FSC curve was calculated from two maps reconstructed from independent two-half datasets. The resolution of the cryoEM maps is depicted in **Fig. S3**.

### Coordinate model building

The coordinate of DENV2 strain PVP94/07 E and M proteins (PDB ID 4UIF) was used to build the coordinate model of DENV2 WT and N153Q mutant, whereas the models of Fabs 290 and C10 (PDB IDs 4UIF and 7V3H) used to fit the density of the respective antibodies.

The coordinate models of E and M proteins and the Fab molecules were first fitted into the cryoEM maps using the “Fit in Map” tool in Chimera program ^47^. The amino acid residues of DENV2 strain PVP94/07 were mutated to the corresponding residues in strain D2Y98P using COOT^48^. The fitted structures were further refined using the “Real-space refinement” tool ^49^ in the Phenix program ^50^ with the secondary structure restraint applied. Several cycles of manual fitting using COOT ^48^ and the real-space refinement were done to achieve good fit parameters. Model refinement and validation statistics are shown in **Tables S1** and **S2**.

### Data availability

The cryoEM maps of DENV2 WT and the N153Q mutant at 4°C, and DENV2 WT – Fab 2D22, DENV2 N153Q – Fab 2D22, DENV2 WT – Fab C10, and DENV2 N153Q – Fab C10 complexes at 37°C were deposited in the Electron Microscopy Database under accession numbers EMD-38884, EMD-38881, EMD-38885, EMD-38882, EMD-38886, and EMD-38883, respectively. The coordinates models of the structures above were deposited in the Protein Data Bank under accession codes 8Y3J, 8Y3G, 8Y3K, 8Y3H, 8Y3L, and 8Y3I, respectively.

### Structure analysis

The atomic displacement parameters were calculated using Phenix real space refinement and the B-factor distribution is displayed in UCSF chimera with tool “Render by Attributes”. The solvent accessible surface areas were calculated by GETAREA webserver (https://curie.utmb.edu/getarea.html) ^51^. The interacting residues on Fab 2D22 and C10, and DENV2 WT and N153Q were identified using the CONTACT from the CCP4 Program Suite^52^ with a threshold distance of 4.2 Å. The Fab occupancy is estimated by the ratio between the average map values at the paratopes residues and the values at the epitope residues (Fab ÷ E protein dimer) x 100 %. All structure figures were prepared with UCSF chimera.

## Acknowledgement

This work was supported by funded by the Ministry of Health (MOH-000087), Singapore awarded to S.M.L and S.A, and the Ministry of Health (MOET32023-0002) awarded to S.M.L.

## Authors contribution

S.M.L. supervised the project. S.A. initiated the project. S.M.L., G.F., and X.-N.L. designed the experiments. J.E.C. and G.R.S. prepared the antibodies. D. H. R. T. and S.A. provided the deglycosylated virus. X.-N.L. prepared the purified deglycosylated viruses and done all the PRNT assays. T.-S.N.,and A.W.K.T. collected the cryoEM images. G.F. performed the cryoEM image processing and reconstruction. G.F. prepared Fabs and performed the BLI experiment. G.F. and S.M.L. interpreted the cryoEM maps, conducted the fitting of molecular models, and structural analysis. G.F., and S.M.L wrote the manuscript.

