## Supplementary information for "Dengue virus 2 lacking N153 glycosylation displayed enhanced recognition by neutralizing antibodies"

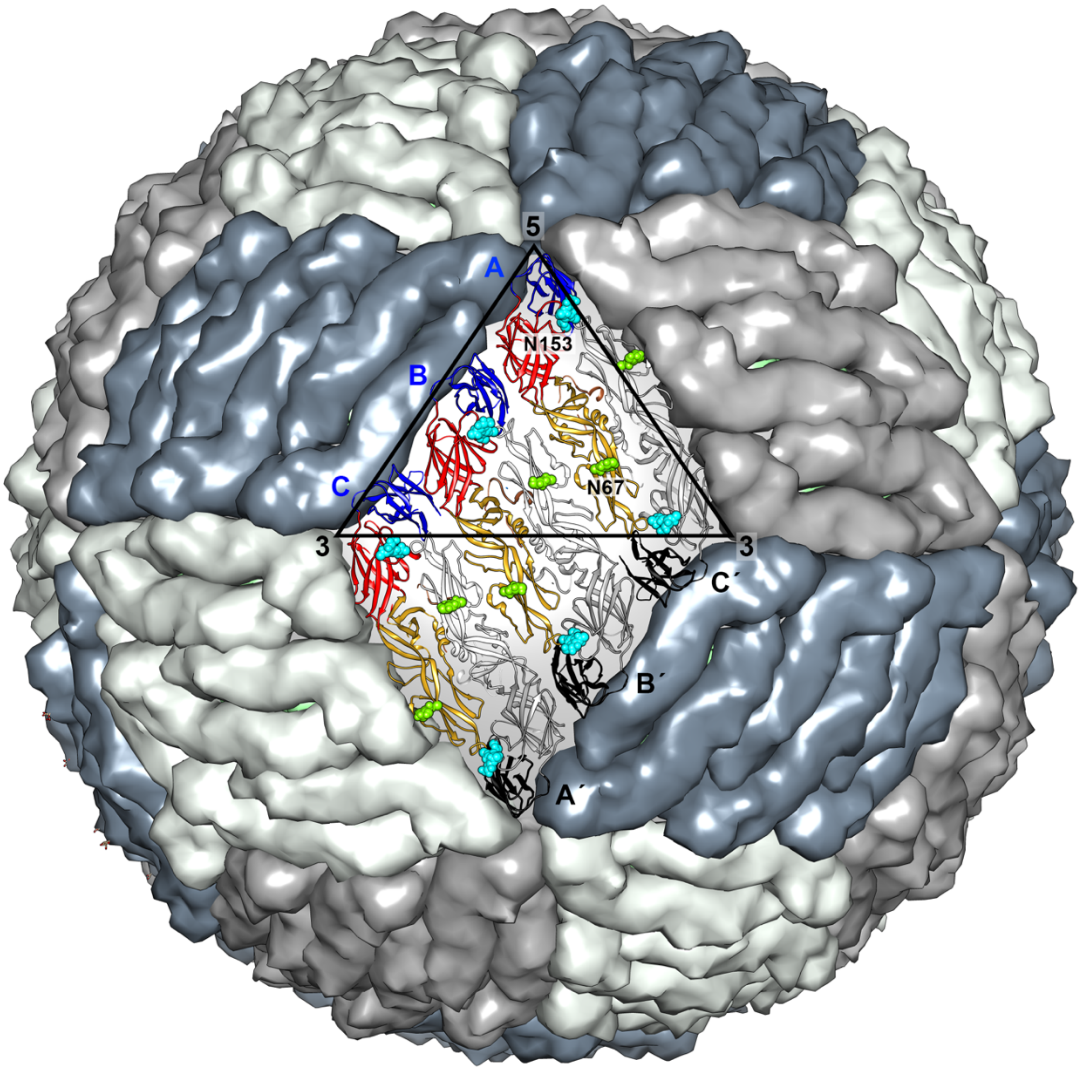

**Figure S1. The arrangement of E proteins on a mature DENV particle.** Each E protein consists of three domains: DI, DII, and DIII. Two E proteins form a head-to-tail dimer, and three of these E protein dimers are arranged parallel to each other, forming a raft structure. A total of 30 E protein rafts are present on the DENV surface, arranged in a herringbone pattern. The E protein rafts are depicted in different shades of grey. One E protein raft is represented in ribbon form, with DI, DII, and DIII of E protein molecules A, B, and C colored in red, yellow, and blue, respectively. The corresponding domains of molecules A´, B´, and C´ are depicted in various shades of grey. Glycosylation at residues N67 and N153 is shown as light green and cyan spheres, respectively. The black triangle indicates an icosahedral asymmetric unit.

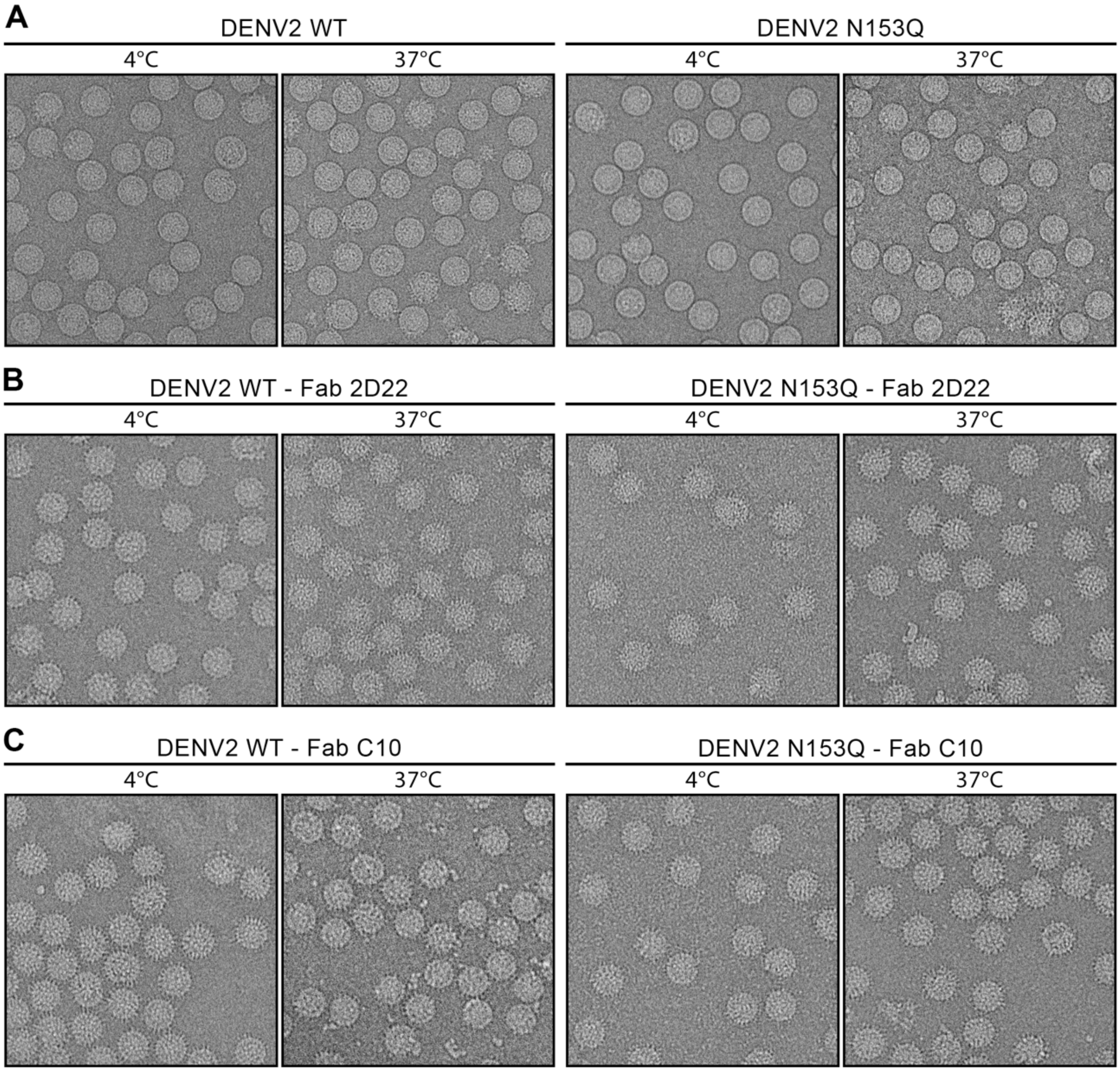

**Figure S2. Cryo-electron micrographs of the uncomplexed DENV2 (WT and N153Q mutant) and the virus complexed with Fab 2D22 or C10 at 4 and 37°C.** Both the DENV2 WT and N153Q mutant maintain their morphology at 37°C. Fabs 2D22 and C10 bind to the DENV2 particles at both 4 and 37°C, resulting in a spiky appearance of the virus particles where the Fabs are bound.

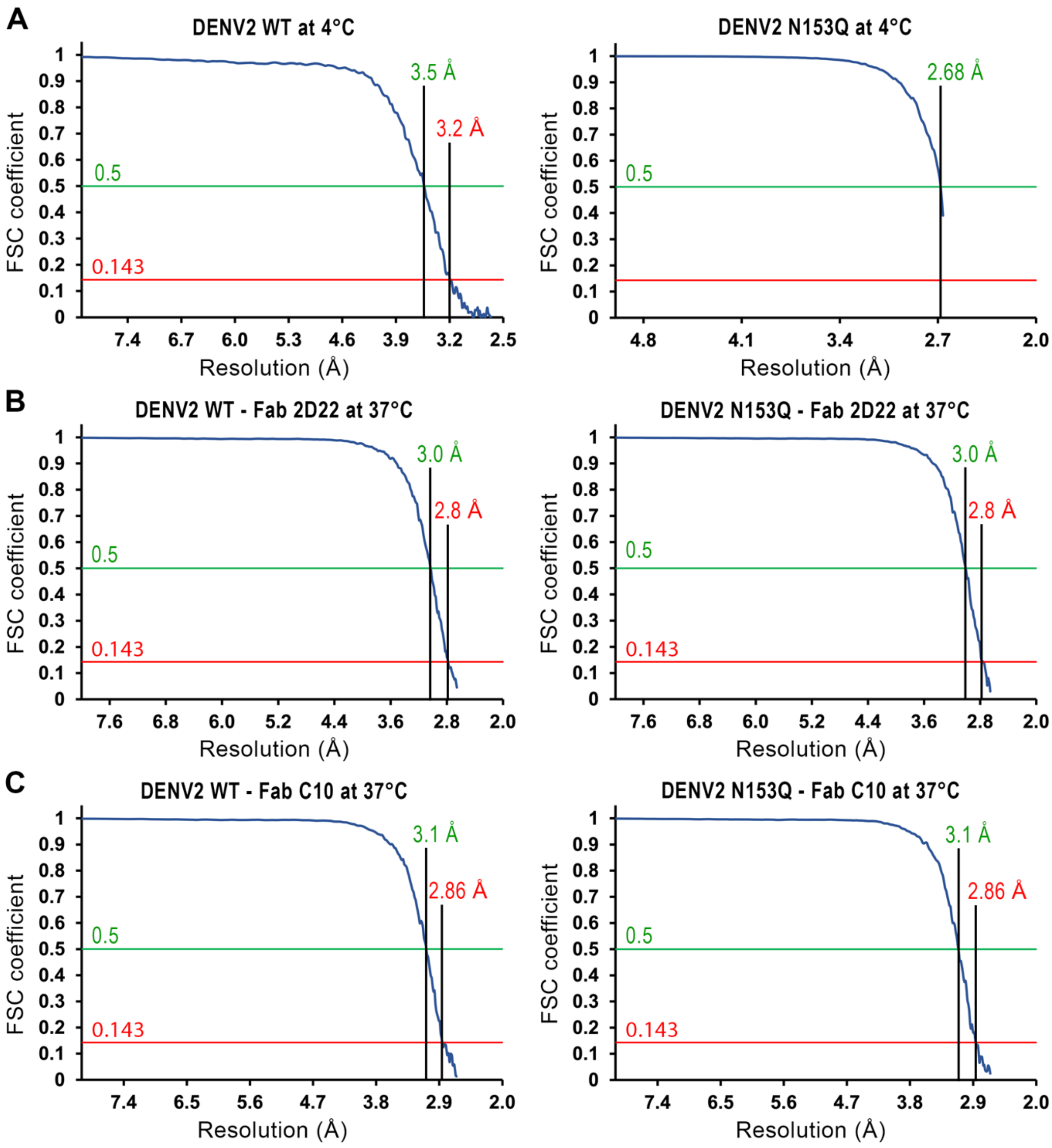

**Figure S3. Resolution assessment of the cryoEM density maps by Fourier Shell Correlation (FSC)**. The FSC curves for all cryoEM maps are displayed. The FSC curve is calculated by comparing two independently reconstructed maps, each derived from a half-dataset, at the final refinement iteration step. The resolutions at the 0.143 and 0.5 FSC cut-off values are indicated.

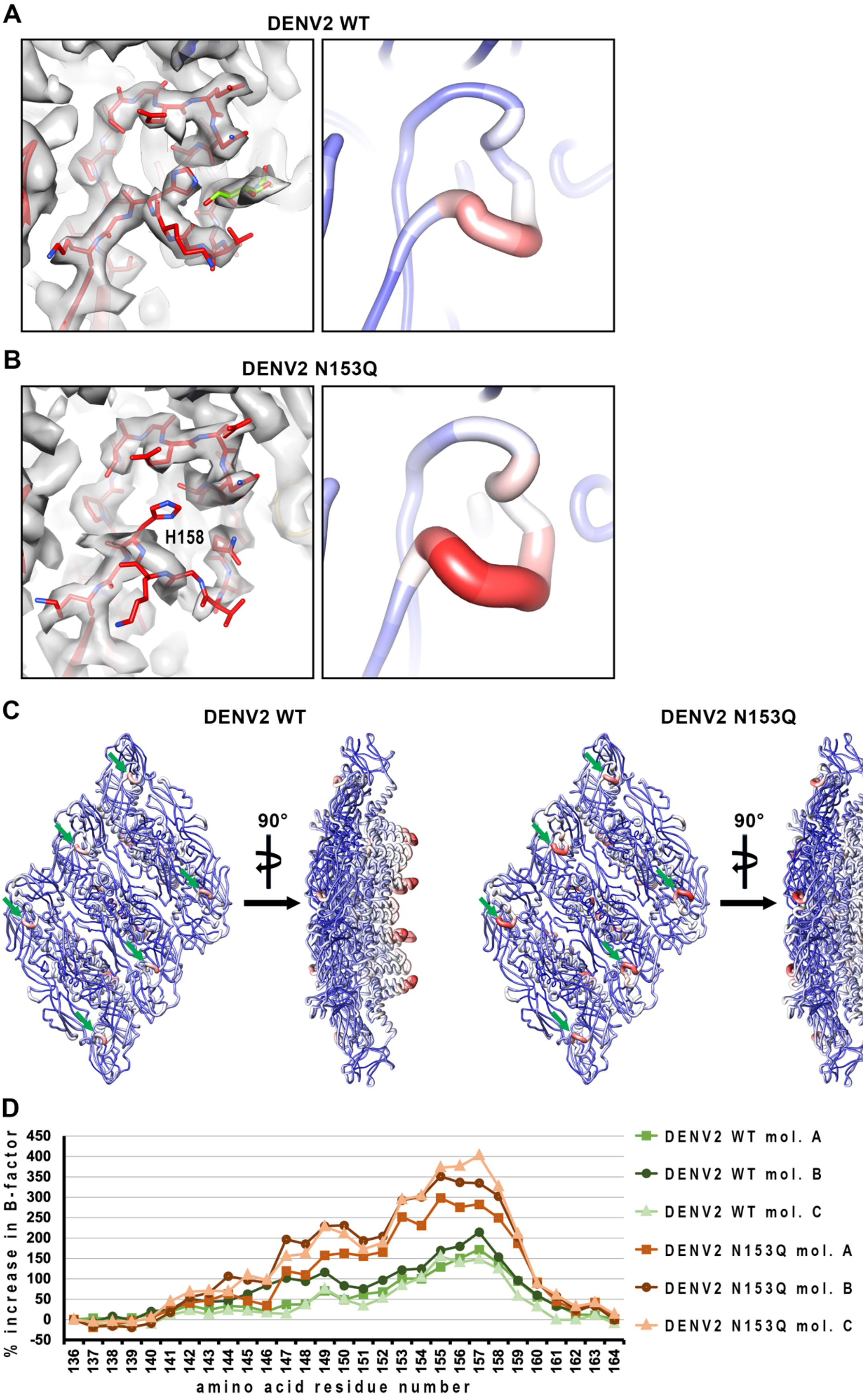
A
DENV2 WT
B
DENV2 N153Q
H158
C
DENV2 WT
90°
DENV2 N153Q
90°
D
% increase in B-factor
450
400
350
300
250
200
150
100
50
0
-50
136 137 138 139 140 141 142 143 144 145 146 147 148 149 150 151 152 153 154 155 156 157 158 159 160 161 162 163 164
amino acid residue number
DENV2 WT mol. A
DENV2 WT mol. B
DENV2 WT mol. C
DENV2 N153Q mol. A
DENV2 N153Q mol. B
DENV2 N153Q mol. C

**Figure S4. The glycan loop in uncomplexed DENV2 N153Q is more dynamic than that in the WT virus.** (**A-B**). [**Left**] The glycan loop densities and the B-factor distribution of the DENV2 WT and N153Q mutant are compared. The glycan loop residues (residues 140-160) of E protein are shown in stick representation, while the surrounding structure is represented as ribbons. The density map is depicted as a transparent grey surface. The glycan loop densities of the DENV2 WT are well defined, while the respective densities in the N153Q mutant are poorer, especially around the residue H158, indicating higher mobility. The glycan residues at N153 are shown in stick representation and colored light green. [**Right**] In the B-factor distribution analysis, the E protein is shown in noodle representation, with color indicating the B-factor values. A larger diameter indicates a higher B-factor. The blue-to-red color gradient also indicates an increasing B-factor, with a value range of 13-84 $Å^2$ for the wild type and 1-53 Å2 for the N153Q mutant. B-factors of protein structures reflect the fluctuation of atoms and provide insight into protein dynamics. Higher B-factors indicate greater protein dynamics. While comparison of B-factors across virus structures is not feasible, as the conditions of the cryoEM data collection environment could contribute to the overall B factors, we can compare within a structure. Comparison of the B-factors of the glycan loops relative to the rest of the E proteins shows the glycan loops in the N153Q mutant have higher mobility, as indicated by a darker red color. (**C**) Overall B-factor distribution of DENV2 WT and N153Q mutant coordinate models is depicted. In both structures, the ectodomain part of the E protein showed lower dynamics, whereas the tip of the transmembrane helices region facing the inner leaflet of the lipid bilayer showed higher dynamics. The glycan loops of all three E proteins in an icosahedral ASU displayed higher mobility (indicated by green arrows). (**D**) B-factor values increase along the glycan loop residues (residues 140-160). The increase in B-factor for each residue is expressed as a percentage relative to the B-factor of residue 136 from the same glycan loop (refer to the equation below). Residue 136 is located on one of the two β-strands flanking the glycan loop and has a relatively low B-factor, similar to that of residues 137-140 or 162-164 on the opposing β-strand.

$$\%\ \text{increase in B} - \text{factor} = \frac{\text{B} - \text{factor (residue)} - \text{B} - \text{factor (residue 136)}}{\text{B} - \text{factor (residue 136)}}\ \text{x}\ 100\%$$

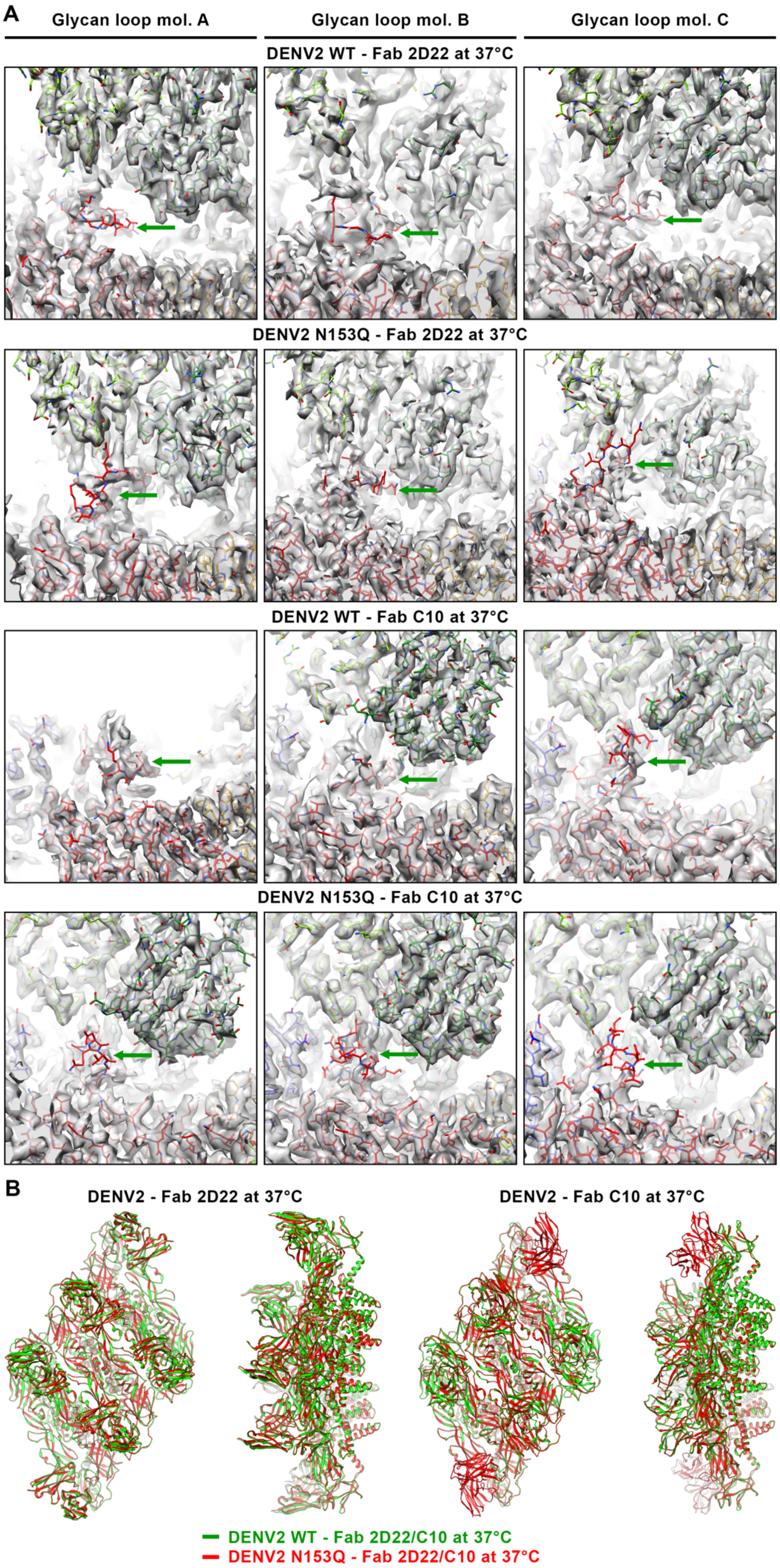
A
Glycan loop mol. A
Glycan loop mol. B
Glycan loop mol. C
DENV2 WT - Fab 2D22 at 37°C
DENV2 N153Q - Fab 2D22 at 37°C
DENV2 WT - Fab C10 at 37°C
DENV2 N153Q - Fab C10 at 37°C
B
DENV2 - Fab 2D22 at 37°C
DENV2 - Fab C10 at 37°C
DENV2 WT - Fab 2D22/C10 at 37°C
DENV2 N153Q - Fab 2D22/C10 at 37°C

**Figure S5. Fabs 2D22 and C10 binding to the E protein disrupted the glycan loop structure.** (**A**) The glycan loop densities became poor when Fab 2D22 or C10 were bound to the DENV2 WT and the N153Q mutant. The coordinate models of the glycan loops (residues 140-160) could not be properly fitted into the densities. Only the glycan loop on E protein molecule A of the DENV2 WT in complex with Fab C10 showed well-defined densities. This is because Fab C10 are likely poorly bound to this site. CryoEM maps are shown as a transparent grey surface, while the E protein and Fab 2D22 and C10 are depicted in stick representation. The E protein DI, DII, and DIII are colored in red, yellow, and blue, respectively, while the heavy and light chains of Fabs 2D22 and C10 are in dark and light green, respectively. The glycan loops on E protein molecules A, B, and C of each virus complex are pointed out with green arrows. (**B**) Superposition of the Fab 2D22 complexed DENV2 WT and N153Q structures and the respective viruses complexed with Fab C10. The superimposed structures of Fabs 2D22 and C10 complexes showed that both Fabs bound to the E protein of the wild type virus and the N153Q mutant in the same binding conformation. There are minimal structural differences between the E proteins of the DENV2 WT and those of the N153Q mutant.

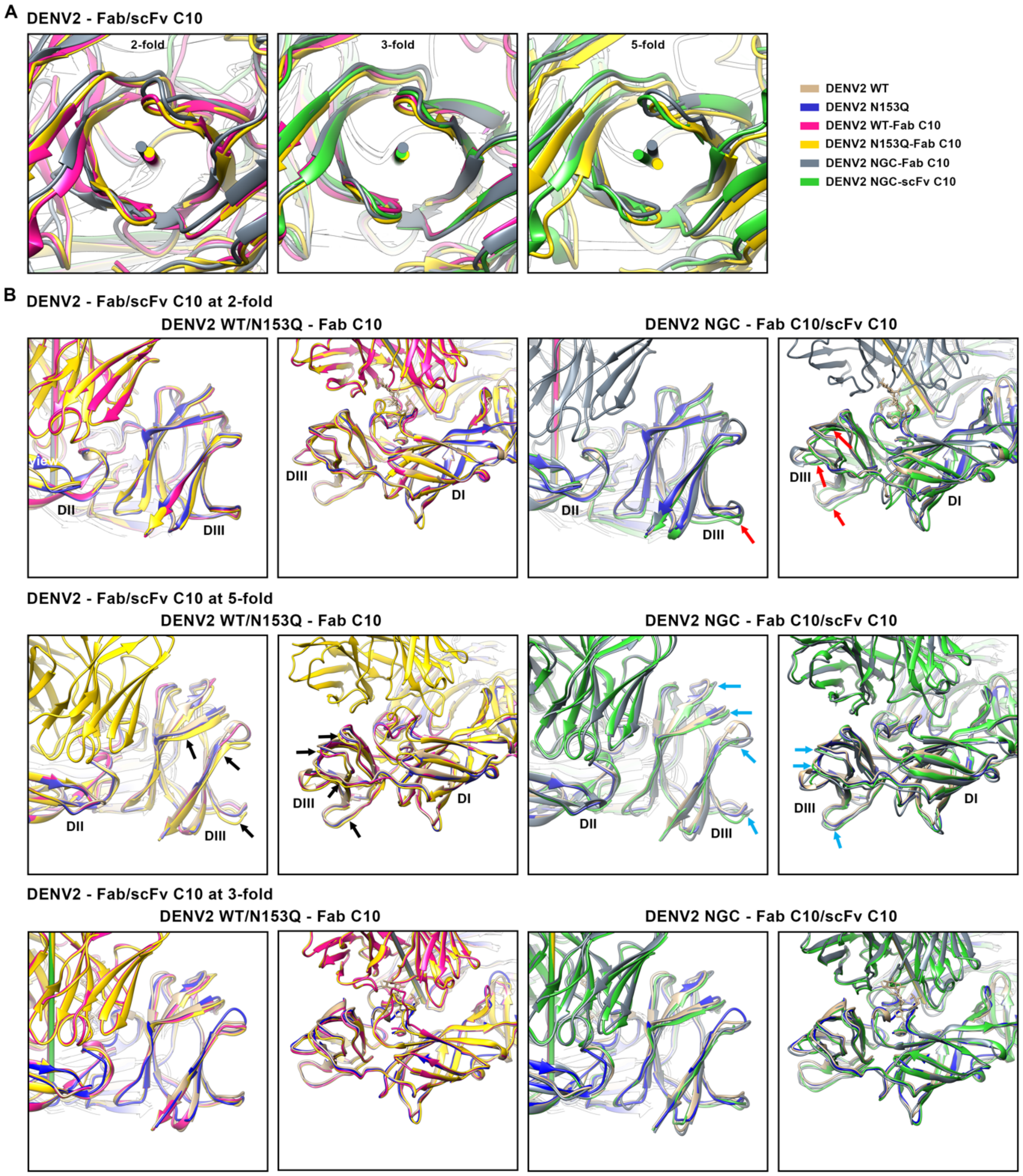

**Figure S6. Conformational changes in the E protein upon binding of Fab C10.** (**A**) The binding axes of Fab C10/scFv C10 on DENV2 WT, N153Q, and DENV2 NGC are similar when Fab C10 binds to epitopes near the 2-fold and 3-fold vertices. However, slight variations are observed when Fab C10 binds near the 5-fold vertices. (**B**) [**Top**, **left**] Superimposition of E proteins at the 2-fold vertices of DENV2 WT/N153Q, with and without Fab C10, showed that the binding of Fab C10 to the E protein near the 2-fold vertices of DENV2 WT or N153Q did not cause any conformational changes in the E proteins. [**Top**, **right**] The structure of the DENV2 NGC – Fab C10 complex (PDB ID: 7V3H) shows that Fab C10 can bind to the E protein at the 2-fold vertices, although with low occupancy (**Fig. 4C**) [1]. However, in the similar structure of the scFv C10 complexed with DENV2 NGC (referred to as NGC-2 here, as it is a substrain with some mutations compared to the DENV2 NGC mentioned earlier, **Fig. S7**; PDB ID: 7CTH), no Fab binding at this location was observed [2]. Superimposition of the E proteins, with and without Fab C10, indicates possible local structural differences at DIII in DENV2 NGC-2 (indicated by red arrows) that affect the Fab C10 binding. For reference, the respective E proteins of uncomplexed DENV2 WT and N153Q, where Fab C10 binds to this epitope, are shown. [**Middle**, **left**] Superimposition of E proteins at the 5-fold vertices of DENV2 WT/N153Q, with and without Fab C10, revealed that the binding of Fab C10 caused conformational changes to the E proteins, particularly in the DIII region (indicated by black arrows). These conformational changes are not observed in the E protein of the DENV2 WT – Fab C10 complex where Fab C10 did not bind to the E protein. [**Middle**, **right**] The structures of DENV2 NGC – Fab C10 and DENV2 NGC2 – scFv C10 complexes showed Fab/scFv C10 binding to the E protein at 5-fold. The superimposition of the E proteins of these complexes at the 5-fold vertices with those of uncomplexed DENV2 WT and N153Q showed differences at DIII region (indicated by blue arrows), similar to what is observed in DENV2 N153Q – Fab C10 complex. [**Bottom**, **left**] Superimposition of E proteins at the 3-fold vertices of DENV2 WT/N153Q, with and without Fab C10, showed that the binding of Fab C10 to the E protein near the 3-fold vertices of DENV2 WT or N153Q did not cause any conformational changes in the E proteins. [**Bottom**, **right**] Similarly, the superimposed structures of the DENV2 NGC in complex with Fab C10 or scFv C10 at the 3-fold vertices show that the Fab/scFv C10 binding does not cause conformational changes when compared to the respective E proteins of uncomplexed DENV2 WT and N153Q. The E protein-Fab complexes, from Fab that binds to epitope near the 2-, 3-, or 5-fold vertices, were aligned separately in the same way as described in **Fig. 5**.

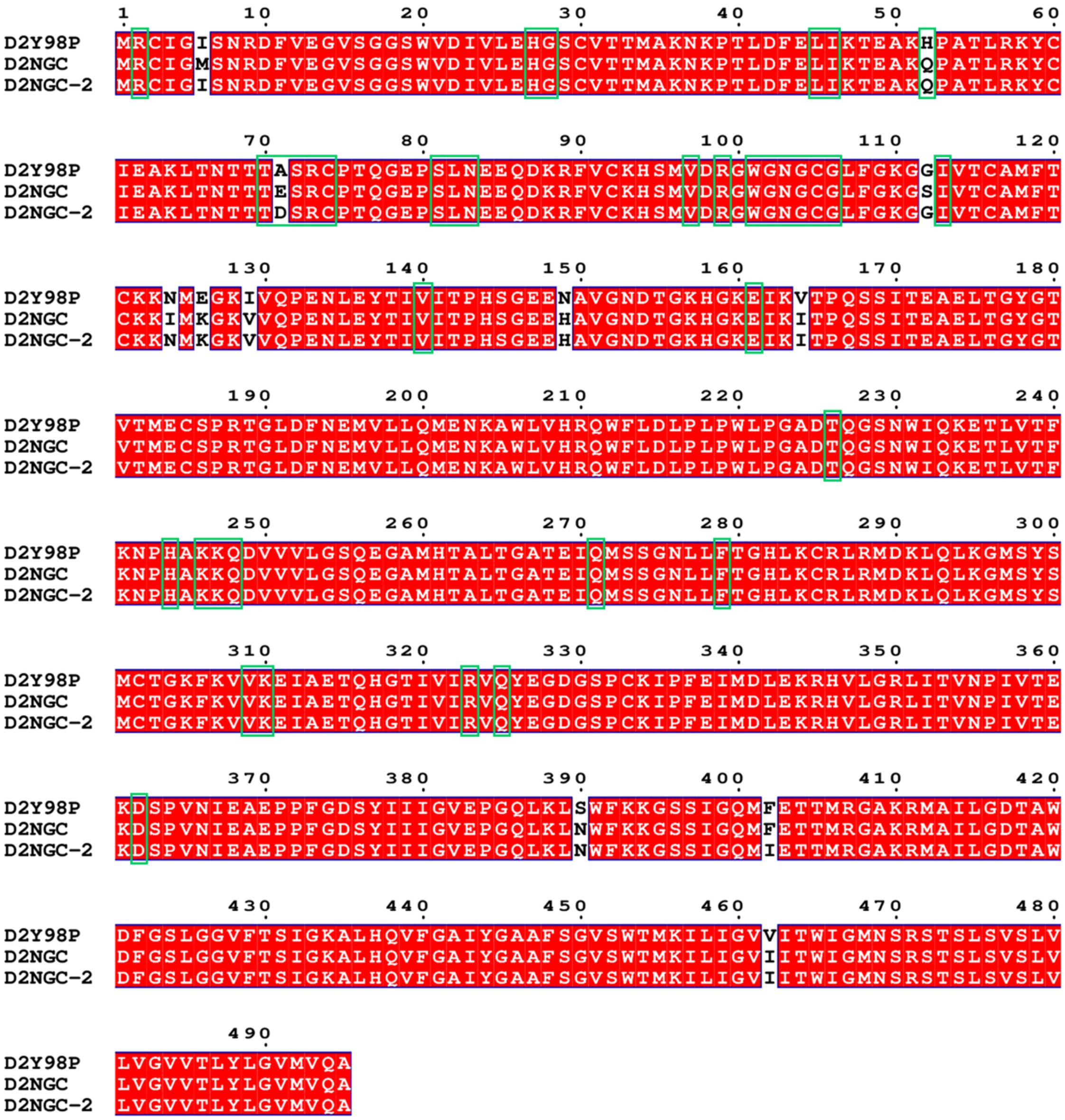

**Figure S7. Sequence alignment of the E protein from DENV2 D2Y98P, DENV2 NGC and DENV2 NGC-2.** DENV2 NGC is the strain used in the structure of the DENV2 NGC – Fab C10 complex, with PDB ID: 7V3H [1]. Whereas, DENV2 NGC-2 is the strain used in the structure of the DENV2 NGC – scFv C10 complex with PDB ID: 7CTH [2].

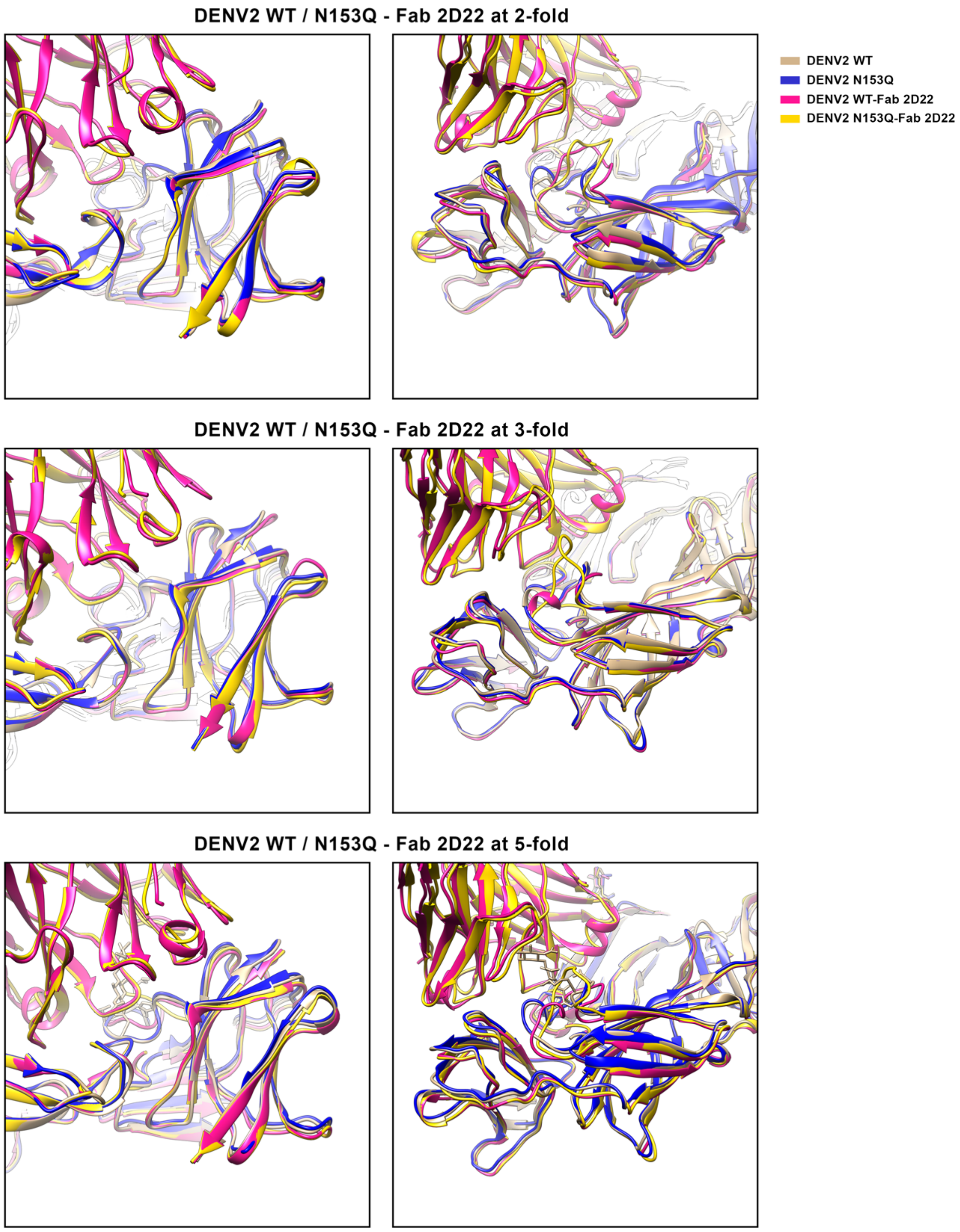

**Figure S8. The binding of Fab 2D22 to DENV2 WT and N153Q mutant does not induce conformational changes in the E protein**. Superposition of E proteins at the 2-, 3-, and 5-fold vertices of DENV2 WT and N153Q mutant, with and without Fab 2D22, showed that the binding of Fab 2D22 to the E protein did not cause significant conformational changes in the

E proteins. The E protein-Fab complexes, from Fabs that bind to epitopes near the 2-, 3-, or 5-fold vertices, were aligned separately in the same way as described in **Fig. 5**.

**Table S1: Cryo-EM data collection, refinement and validation statistics of DENV2 WT and N153Q**

| | ***DENV2 Y98P-PP1 at 4 °C*** (EMD-38884, PDB 8Y3J) | **DENV2 Y98P-PP1 N153Q at 4 °C** (EMD-38881, PDB 8Y3G) |
|---|---|---|
| **Data collection and processing** | | |
| Microscope | FEI Titan Krios | FEI Titan Krios |
| Voltage (kV) | 300 | 300 |
| Camera | Gatan K3 | Gatan K3 |
| Pixel size (Å) | 1.345 | 1.345 |
| Magnification | 64,000 | 64,000 |
| Number of micrographs | 1,980 | 8,533 |
| Dose rate e-/Å$^2$.s$^{-1}$ | 5.27 | 5.78 |
| Frames per movie | 18 | 25 |
| Total electron exposure (e$^-$/Å$^2$) | 23.75 | 25 |
| Defocus range (μm) | -0.4 to -5.2 | -0.4 to -5.1 |
| Symmetry imposed | Icosahedral | Icosahedral |
| Initial particle images (no.) | 212,371 | 640,232 |
| Final particle images (no.) | 14,421 | 173,676 |
| Map-sharpening B-factor (Å$^2$) | -100 | -94 |
| Map Resolution (Å) | 3.2 | 2.7 |
| FSC threshold | 0.143 | 0.143 |
| **Model refinement** | | |
| Initial model used (PDB code) | 4UIF | 4UIF |
| Model resolution at 0.5 FSC threshold (Å) | 3.2 | 2.7 |
| Model composition | | |
| Non-hydrogen atoms | 14,910 | 14,829 |
| Protein residues | 1,710 | 1,710 |
| Ligand | 9 | 3 |
| Global CC (CC volume) | 0.81 | 0.78 |
| Local CC (CC mask) | 0.87 | 0.87 |
| B factors (Å$^2$) | | |
| Protein | 31.31 | 14.75 |
| Ligand | 66.82 | 30.43 |
| R.M.S.D. deviations | | |
| Bond lengths (Å) | 0.005 | 0.003 |
| Bond angles (°) | 0.527 | 0.497 |
| **Validation** | | |
| MolProbity score | 2.05 | 2.27 |
| Clashscore | 25.49 | 25.25 |
| Rotamers outliers (%) | 0.00 | 0.00 |
| Ramachandran plot | | |
| Favored (%) | 97.17 | 94.46 |
| Allowed (%) | 2.83 | 5.54 |
| Disallowed (%) | 0.00 | 0.00 |
| CaBLAM outliers (%) | 2.14 | 4.09 |

**Table S2: Cryo-EM data collection, refinement and validation statistics of DENV2 WT and N153Q complexes**

| | **DENV2 Y98P-PP1 - Fab 2D22 at 37 ºC** (EMD-38885, PDB 8Y3K) | **DENV2 Y98P-PP1 N153Q - Fab 2D22 at 37 ºC** (EMD-38882, PDB 8Y3H) | **DENV2 Y98P-PP1 - Fab C10 at 37 ºC** (EMD-38886, PDB 8Y3L) | **DENV2 Y98P-PP1 N153Q - Fab C10 at 37 ºC** (EMD-38883, PDB 8Y3I) |
|---|---|---|---|---|
| **Data collection and processing** | | | | |
| Microscope | FEI Titan Krios | FEI Titan Krios | FEI Titan Krios | FEI Titan Krios |
| Voltage (kV) | 300 | 300 | 300 | 300 |
| Camera | Gatan K3 | Gatan K3 | Gatan K3 | Gatan K3 |
| Pixel size (Å) | 1.345 | 1.345 | 1.345 | 1.345 |
| Magnification | 64,000 | 64,000 | 64,000 | 64,000 |
| Number of micrographs | 4,778 | 4,919 | 5,921 | 4,439 |
| Dose rate $e^-/Å^2.s^{-1}$ | 5.40 | 5.15 | 5.42 | 5.19 |
| Frames per movie | 19 | 18 | 18 | 18 |
| Total electron exposure ($e^-/Å^{-2}$) | 25.69 | 23.22 | 24.43 | 23.37 |
| Defocus range (μm) | -0.5 to -4.4 | -0.5 to -5.0 | -0.5 to -3.7 | -0.5 to -4.7 |
| Symmetry imposed | Icosahedral | Icosahedral | Icosahedral | Icosahedral |
| Initial particle images (no.) | 281,127 | 272,647 | 337,394 | 242,187 |
| Final particle images (no.) | 123,807 | 149,073 | 64,522 | 102,447 |
| Map-sharpening B-factor ($Å^2$) | -107 | -106 | -104 | -119 |
| Map Resolution (Å) | 2.8 | 2.8 | 2.9 | 2.9 |
| FSC threshold | 0.143 | 0.143 | 0.143 | 0.143 |
| **Model refinement** | | | | |
| Initial model used (PDB code) | 4UIF | 4UIF | 4UIF, 5H37 | 4UIF, 5H37 |
| Model resolution at 0.5 FSC threshold (Å) | 2.8 | 2.9 | 2.9 | 2.9 |
| Model composition | | | | |
| Non-hydrogen atoms | 20,042 | 20,049 | 18,471 | 20,283 |
| Protein residues | 2,403 | 2,403 | 2,182 | 2,419 |
| Ligand | 3 | 3 | 4 | 3 |
| Global CC (CC volume) | 0.74 | 0.74 | 0.76 | 0.74 |
| Local CC (CC mask) | 0.83 | 0.83 | 0.81 | 0.82 |
| B factors ($Å^2$) | | | | |
| Protein | 11.20 | 15.76 | 195.60 | 17.91 |
| Ligand | 20.79 | 21.59 | 29.02 | 32.25 |
| R.M.S.D. deviations | | | | |
| Bond lengths (Å) | 0.005 | 0.003 | 0.004 | 0.004 |
| Bond angles (°) | 0.542 | 0.487 | 0.499 | 0.623 |
| **Validation** | | | | |
| MolProbity score | 2.20 | 2.27 | 2.22 | 2.27 |
| Clashscore | 27.23 | 30.36 | 27.76 | 32.36 |
| Rotamers outliers (%) | 0.10 | 0 | 0 | 0.00 |
| Ramachandran plot | | | | |
| Favored (%) | 95.96 | 95.59 | 95.79 | 96.03 |
| Allowed (%) | 4.04 | 4.41 | 4.16 | 3.97 |
| Disallowed (%) | 0.00 | 0.00 | 0.05 | 0.00 |
| CaBLAM outliers | 2.85 | 3.23 | 2.52 | 2.19 |

**Table S3. Interacting residues between DENV2 WT and Fab 2D22 at 37ºC (Fab 2D22 binds to an epitope located near the 5-fold vertices)**

| No. | Fab 2D22 chain H | mol. A | mol. C' | mol. A* | mol. B* | Fab 2D22 chain L | mol. A | mol. C' | mol. A* | mol. B* |
|---|---|---|---|---|---|---|---|---|---|---|
| 1 | S17 | | | | D329 | S26 | | | S298 | |
| 2 | N30 | | K247 | | | | | | Y299 | |
| 3 | I52 | | N103 | | | S30 | K307 | | | |
| 4 | I54 | | T70 | | | | Q325 | | | |
| 5 | | | K247 | | | S31 | | | S298 | |
| 6 | F55 | | T70 | | | | V309 | | | |
| 7 | | | A71 | | | Y33 | R323 | | | |
| 8 | | | S72 | | | | P364 | | | |
| 9 | | | V97 | | | N52 | D362 | | | |
| 10 | | | R99 | | | N53 | E360 | | | |
| 11 | | | N103 | | | | D362 | | | |
| 12 | | | K246 | | | K67 | D362 | | | |
| 13 | G56 | | T70 | | | D94 | | | Q293 | |
| 14 | | | S72 | | | | V309 | | | |
| 15 | G57 | | S72 | | | S95 | | | E174 | |
| 16 | Q62 | | | E174 | | | | | G177 | |
| 17 | | | | T180 | | | | | Y178 | |
| 18 | G66 | | | | K305 | | | | G179 | |
| 19 | | | | | G385 | | | | Q293 | |
| 20 | R67 | | | | K305 | | | | | |
| 21 | V68 | | | | P384 | | | | | |
| 22 | T69 | | | | P384 | | | | | |
| 23 | A72 | | T69 | | | | | | | |
| 24 | D73 | | N67 | | | | | | | |
| 25 | | | T68 | | | | | | | |
| 26 | R74 | | T68 | | | | | | | |
| 27 | | | T70 | | | | | | | |
| 28 | | | K247 | | | | | | | |
| 29 | | | Q248 | | | | | | | |
| 30 | | | D249 | | | | | | | |
| 31 | S75 | | T66 | | | | | | | |
| 32 | | | N67 | | | | | | | |
| 33 | | | T68 | | | | | | | |
| 34 | S84 | | | | K305 | | | | | |
| 35 | Q101 | | G102 | | | | | | | |
| 36 | | | N103 | | | | | | | |
| 37 | | | G104 | | | | | | | |
| 38 | S102 | | W101 | | | | | | | |
| 39 | | | G102 | | | | | | | |
| 40 | I103 | | W101 | | | | | | | |
| 41 | | | G104 | | | | | | | |
| 42 | | | C105 | | | | | | | |
| 43 | | | G106 | | | | | | | |
| 44 | | K310 | | | | | | | | |
| 45 | F104 | | W101 | | | | | | | |
| 46 | | K310 | | | | | | | | |
| 47 | | R323 | | | | | | | | |

**Table S4. Interacting residues between DENV2 WT and Fab 2D22 at 37ºC (Fab 2D22 binds to an epitope located near the 2-fold vertices)**

| No. | Fab 2D22 chain H | mol. B | mol. B' | mol. A | Fab 2D22 chain L | mol. B | mol. B' | mol. A |
|---|---|---|---|---|---|---|---|---|
| 1 | N30 | | K247 | | S26 | K307 | | |
| 2 | I52 | | N103 | | G30 | K307 | | |
| 3 | P53 | | T70 | | | Q325 | | |
| 4 | I54 | | T70 | | S31 | K307 | | |
| 5 | | | I113 | | | V309 | | |
| 6 | F55 | | T70 | | Y33 | R323 | | |
| 7 | | | A71 | | | P364 | | |
| 8 | | | S72 | | N52 | D362 | | |
| 9 | | | V97 | | N53 | E360 | | |
| 10 | | | R99 | | | D362 | | |
| 11 | | | N103 | | K67 | D362 | | |
| 12 | | | I113 | | D94 | V309 | | |
| 13 | | | K246 | | | | | |
| 14 | G56 | | T70 | | | | | |
| 15 | | | A71 | | | | | |
| 16 | | | S72 | | | | | |
| 17 | G57 | | A71 | | | | | |
| 18 | | | S72 | | | | | |
| 19 | A72 | | T68 | | | | | |
| 20 | | | T69 | | | | | |
| 21 | | | T70 | | | | | |
| 22 | D73 | | N67 | | | | | |
| 23 | | | T68 | | | | | |
| 24 | | | T69 | | | | | |
| 25 | | | glycan | | | | | |
| 26 | R74 | | T68 | | | | | |
| 27 | | | T70 | | | | | |
| 28 | | | T115 | | | | | |
| 29 | | | K247 | | | | | |
| 30 | | | Q248 | | | | | |
| 31 | S75 | | T66 | | | | | |
| 32 | | | N67 | | | | | |
| 33 | | | T68 | | | | | |
| 34 | Y80 | | glycan | | | | | |
| 35 | Q101 | | G102 | | | | | |
| 36 | | | N103 | | | | | |
| 37 | | | G104 | | | | | |
| 38 | S102 | | W101 | | | | | |
| 39 | | | G102 | | | | | |
| 40 | I103 | | W101 | | | | | |
| 41 | | | G104 | | | | | |
| 42 | | | C105 | | | | | |
| 43 | F104 | | W101 | | | | | |
| 44 | | K310 | | | | | | |
| 45 | | R323 | | | | | | |

**Table S5. Interacting residues between DENV2 WT and Fab 2D22 at 37ºC (Fab 2D22 binds to an epitope located near the 3-fold vertices)**

| No. | Fab 2D22 chain H | mol. C | mol. A' | mol. B | Fab 2D22 chain L | mol. C | mol. A' | mol. B |
|---|---|---|---|---|---|---|---|---|
| 1 | N30 | | K247 | | G30 | K307 | | |
| 2 | I52 | | N103 | | | V309 | | |
| 3 | | | N103 | | | Q325 | | |
| 4 | I54 | | T70 | | S31 | V309 | | |
| 5 | | | K247 | | Y33 | R323 | | |
| 6 | F55 | | T70 | | | P364 | | |
| 7 | | | A71 | | N52 | D362 | | |
| 8 | | | S72 | | N53 | D362 | | |
| 9 | | | V97 | | K67 | Q325 | | |
| 10 | | | R99 | | | D362 | | |
| 11 | | | N103 | | G69 | K307 | | |
| 12 | | | I113 | | D94 | V309 | | |
| 13 | | | K246 | | S95 | | | H52 |
| 14 | G56 | | T70 | | | | | |
| 15 | | | A71 | | | | | |
| 16 | | | S72 | | | | | |
| 17 | G57 | | A71 | | | | | |
| 18 | | | S72 | | | | | |
| 19 | A72 | | T68 | | | | | |
| 20 | | | T69 | | | | | |
| 21 | | | T70 | | | | | |
| 22 | D73 | | N67 | | | | | |
| 23 | | | T68 | | | | | |
| 24 | | | glycan | | | | | |
| 25 | R74 | | T68 | | | | | |
| 26 | | | T70 | | | | | |
| 27 | | | T115 | | | | | |
| 28 | | | K247 | | | | | |
| 29 | | | Q248 | | | | | |
| 30 | | | D249 | | | | | |
| 31 | S75 | | T66 | | | | | |
| 32 | | | N67 | | | | | |
| 33 | | | T68 | | | | | |
| 34 | T76 | | glycan | | | | | |
| 35 | Q101 | | G102 | | | | | |
| 36 | | | N103 | | | | | |
| 37 | | | G104 | | | | | |
| 38 | S102 | | W101 | | | | | |
| 39 | | | G102 | | | | | |
| 40 | I103 | | W101 | | | | | |
| 41 | | | G104 | | | | | |
| 42 | | | C105 | | | | | |
| 43 | | K310 | | | | | | |
| 44 | F104 | | W101 | | | | | |
| 45 | | K310 | | | | | | |
| 46 | | R323 | | | | | | |

**Table S6. Interacting residues between DENV2 WT and Fab C10 at 37ºC (Fab C10 binds to an epitope located near to 2-fold vertices)**

| No. | Fab C10 chain H | mol. B | mol. B' | mol. A | Fab C10 chain L | mol. B | mol. B' | mol. A |
|---|---|---|---|---|---|---|---|---|
| 1 | Q65 | | glycan | | G31 | | C74 | |
| 2 | D103 | R2 | | | F32 | | S72 | |
| 3 | | I46 | | | | | R73 | |
| 4 | | E161 | | | | | C74 | |
| 5 | Y104 | H27 | | | | | R99 | |
| 6 | | G28 | | | | | G104 | |
| 7 | | L45 | | | | | C105 | |
| 8 | | I46 | | | N33 | | G104 | |
| 9 | | Q271 | | | | | C105 | |
| 10 | | F279 | | | | | G106 | |
| 11 | | | H244 | | N33 | K310 | | |
| 12 | D106 | | K247 | | Y34 | | G102 | |
| 13 | W108 | | T70 | | | | N103 | |
| 14 | | | I113 | | | | G104 | |
| 15 | | | K247 | | D52 | R323 | | |
| 16 | | | Q248 | | S55 | R323 | | |
| 17 | F109 | | S72 | | R56 | D362 | | |
| 18 | | | R99 | | S62 | D362 | | |
| 19 | | | N103 | | S69 | | | H52 |
| 20 | L112 | | W101 | | G70 | | | H52 |
| 21 | | | G102 | | S95 | | T70 | |
| 22 | | | | | | | A71 | |
| 23 | | | | | | | S72 | |
| 24 | | | | | R96 | | T70 | |

**Table S7. Interacting residues between DENV2 WT and Fab C10 at 37ºC (Fab C10 binds to an epitope located near to 3-fold vertices)**

| No. | Fab C10 chain H | mol. C | mol. A' | mol. B | Fab C10 chain L | mol. C | mol. A' | mol. B |
|---|---|---|---|---|---|---|---|---|
| 1 | D102 | | K246 | | S26 | | R73 | |
| 2 | D103 | R2 | | | G31 | | C74 | |
| 3 | | I46 | | | F32 | | S72 | |
| 4 | | V140 | | | | | R73 | |
| 5 | Y104 | H27 | | | | | C74 | |
| 6 | | | H244 | | | | R99 | |
| 7 | | | K246 | | | | G104 | |
| 8 | | F279 | | | | | C105 | |
| 9 | D106 | | K247 | | N33 | | G104 | |
| 10 | W108 | | V97 | | | | C105 | |
| 11 | | | I113 | | | | G106 | |
| 12 | | | K246 | | | K310 | | |
| 13 | | | K247 | | Y34 | | N103 | |
| 14 | F109 | | S72 | | | | G104 | |
| 15 | | | R99 | | D52 | K310 | | |
| 16 | | | N103 | | | R323 | | |
| 17 | L112 | | W101 | | T54 | V309 | | |
| 18 | | | G102 | | S55 | R323 | | |
| 19 | | | G104 | | R56 | Q325 | | |
| 20 | | | | | | D362 | | |
| 21 | | | | | S62 | D362 | | |
| 22 | | | | | S69 | | | H52 |
| 23 | | | | | G70 | | | H52 |
| 24 | | | | | S95 | | T70 | |
| 25 | | | | | | | A71 | |
| 26 | | | | | | | S72 | |
| 27 | | | | | R96 | | T70 | |
| 28 | | | | | | | A71 | |
| 29 | | | | | | | S81 | |
| 30 | | | | | | | L82 | |
| 31 | | | | | | | N83 | |
| 32 | | | | | | | | T226 |

**Table S8. Interacting residues between DENV2 N153Q and Fab 2D22 at 37ºC (Fab 2D22 binds to an epitope located near to 5-fold vertices)**

| No. | Fab 2D22 chain H | mol. A | mol. C' | mol. A* | mol. B* | Fab 2D22 chain L | mol. A | mol. C' | mol. A* | mol. B* |
|---|---|---|---|---|---|---|---|---|---|---|
| 1 | S17 | | | | D329 | S26 | | | S298 | |
| 2 | N30 | | K247 | | | | | | Y299 | |
| 3 | I52 | | R99 | | | | K307 | | | |
| 4 | | | N103 | | | G30 | K307 | | | |
| 5 | P53 | | T70 | | | | Q325 | | | |
| 6 | I54 | | T70 | | | S31 | | | S298 | |
| 7 | | | K247 | | | | V309 | | | |
| 8 | F55 | | T70 | | | | Q325 | | | |
| 9 | | | A71 | | | Y33 | R323 | | | |
| 10 | | | S72 | | | | P364 | | | |
| 11 | | | V97 | | | N52 | D362 | | | |
| 12 | | | R99 | | | N53 | E360 | | | |
| 13 | | | N103 | | | K67 | Q325 | | | |
| 14 | | | I113 | | | | D362 | | | |
| 15 | | | K246 | | | S68 | D362 | | | |
| 16 | G56 | | T70 | | | D94 | | | Q293 | |
| 17 | | | S72 | | | | V309 | | | |
| 18 | G57 | | S72 | | | S95 | | | G177 | |
| 19 | Q62 | | | E174 | | | | | G179 | |
| 20 | G66 | | | | K305 | | | | Q293 | |
| 21 | | | | | P384 | | | | | |
| 22 | T69 | | | | P384 | | | | | |
| 23 | A72 | | T68 | | | | | | | |
| 24 | D73 | | N67 | | | | | | | |
| 25 | | | T68 | | | | | | | |
| 26 | R74 | | T68 | | | | | | | |
| 27 | | | T70 | | | | | | | |
| 28 | | | K247 | | | | | | | |
| 29 | | | Q248 | | | | | | | |
| 30 | | | D249 | | | | | | | |
| 31 | S75 | | T66 | | | | | | | |
| 32 | | | N67 | | | | | | | |
| 33 | | | T68 | | | | | | | |
| 34 | S84 | | | | K305 | | | | | |
| 35 | Q101 | | W101 | | | | | | | |
| 36 | | | G102 | | | | | | | |
| 37 | | | N103 | | | | | | | |
| 38 | | | G104 | | | | | | | |
| 39 | S102 | | W101 | | | | | | | |
| 40 | | | G102 | | | | | | | |
| 41 | I103 | | W101 | | | | | | | |
| 42 | | | G104 | | | | | | | |
| 43 | | | C105 | | | | | | | |
| 44 | | | G106 | | | | | | | |
| 45 | | K310 | | | | | | | | |
| 46 | F104 | M1 | | | | | | | | |
| 47 | | | W101 | | | | | | | |
| 48 | | K310 | | | | | | | | |
| 49 | | R323 | | | | | | | | |

**Table S9. Interacting residues between DENV2 N153Q and Fab 2D22 at 37ºC (Fab 2D22 binds to an epitope located near to 2-fold vertices)**

| No. | Fab 2D22 chain H | mol. B | mol. B' | mol. A | Fab 2D22 chain L | mol. B | mol. B' | mol. A |
|---|---|---|---|---|---|---|---|---|
| 1 | N30 | | K247 | | G30 | K307 | | |
| 2 | I52 | | N103 | | | Q325 | | |
| 3 | | | G104 | | S31 | K307 | | |
| 4 | P53 | | T70 | | | V309 | | |
| 5 | I54 | | T70 | | | Q325 | | |
| 6 | | | I113 | | Y33 | R323 | | |
| 7 | | | K247 | | | P364 | | |
| 8 | F55 | | T70 | | R51 | E360 | | |
| 9 | | | A71 | | N52 | D362 | | |
| 10 | | | S72 | | N53 | E360 | | |
| 11 | | | V97 | | | D362 | | |
| 12 | | | R99 | | K67 | D362 | | |
| 13 | | | N103 | | G69 | K307 | | |
| 14 | | | K246 | | D94 | V309 | | |
| 15 | G56 | | T70 | | | | | |
| 16 | | | A71 | | | | | |
| 17 | | | S72 | | | | | |
| 18 | G57 | | A71 | | | | | |
| 19 | | | S72 | | | | | |
| 20 | T69 | | N83 | | | | | |
| 21 | A72 | | T68 | | | | | |
| 22 | | | T69 | | | | | |
| 23 | D73 | | N67 | | | | | |
| 24 | | | T68 | | | | | |
| 25 | | | glycan | | | | | |
| 26 | R74 | | T68 | | | | | |
| 27 | | | T70 | | | | | |
| 28 | | | T115 | | | | | |
| 29 | | | K247 | | | | | |
| 30 | | | Q248 | | | | | |
| 31 | | | D249 | | | | | |
| 32 | S75 | | T66 | | | | | |
| 33 | | | N67 | | | | | |
| 34 | | | T68 | | | | | |
| 35 | Q101 | | G102 | | | | | |
| 36 | | | N103 | | | | | |
| 37 | | | G104 | | | | | |
| 38 | S102 | | W101 | | | | | |
| 39 | | | G102 | | | | | |
| 40 | | | G104 | | | | | |
| 41 | I103 | | W101 | | | | | |
| 42 | | | G104 | | | | | |
| 43 | | | C105 | | | | | |
| 44 | | | G106 | | | | | |
| 45 | F104 | | W101 | | | | | |
| 46 | | K310 | | | | | | |
| 47 | | R323 | | | | | | |

**Table S10. Interacting residues between DENV2 N153Q and Fab 2D22 at 37ºC (Fab 2D22 binds to an epitope located near to 3-fold vertices)**

| No. | Fab 2D22 chain H | mol. C | mol. A' | mol. B | Fab 2D22 chain L | mol. C | mol. A' | mol. B |
|---|---|---|---|---|---|---|---|---|
| 1 | N30 | | K247 | | G30 | K307 | | |
| 2 | I52 | | N103 | | | V309 | | |
| 3 | I54 | | T70 | | | Q325 | | |
| 4 | | | K247 | | S31 | K307 | | |
| 5 | F55 | | T70 | | | V309 | | |
| 6 | | | A71 | | Y33 | R323 | | |
| 7 | | | S72 | | | P364 | | |
| 8 | | | V97 | | N52 | D362 | | |
| 9 | | | R99 | | K67 | D362 | | |
| 10 | | | N103 | | S68 | D362 | | |
| 11 | | | I113 | | | | | |
| 12 | | | K246 | | | | | |
| 13 | G56 | | T70 | | | | | |
| 14 | | | A71 | | | | | |
| 15 | | | S72 | | | | | |
| 16 | G57 | | A71 | | | | | |
| 17 | | | S72 | | | | | |
| 18 | A72 | | T69 | | | | | |
| 19 | D73 | | N67 | | | | | |
| 20 | | | T68 | | | | | |
| 21 | | | glycan | | | | | |
| 22 | R74 | | T68 | | | | | |
| 23 | | | T70 | | | | | |
| 24 | | | T115 | | | | | |
| 25 | | | K247 | | | | | |
| 26 | | | Q248 | | | | | |
| 27 | | | D249 | | | | | |
| 28 | S75 | | T66 | | | | | |
| 29 | | | N67 | | | | | |
| 30 | | | T68 | | | | | |
| 31 | T76 | | glycan | | | | | |
| 32 | Q101 | | G102 | | | | | |
| 33 | | | N103 | | | | | |
| 34 | | | G104 | | | | | |
| 35 | S102 | M1 | | | | | | |
| 36 | | | W101 | | | | | |
| 37 | | | G102 | | | | | |
| 38 | I103 | | W101 | | | | | |
| 39 | | | G104 | | | | | |
| 40 | | | C105 | | | | | |
| 41 | | | G106 | | | | | |
| 42 | | K310 | | | | | | |
| 43 | F104 | M1 | | | | | | |
| 44 | | | W101 | | | | | |
| 45 | | K310 | | | | | | |
| 46 | | R323 | | | | | | |

**Table S11. Interacting residues between DENV2 N153Q and Fab C10 at 37ºC (Fab C10 binds to an epitope located near to 5-fold vertices)**

| No. | Fab C10 chain H | mol. A | mol. C' | mol. A* | mol. B* | Fab C10 chain L | mol. A | mol. C' | mol. A* | mol. B* |
|---|---|---|---|---|---|---|---|---|---|---|
| 1 | Q65 | | glycan | | | S26 | | | K291 | |
| 2 | D103 | R2 | | | | G30 | | | K291 | |
| 3 | | I46 | | | | G31 | | C74 | | |
| 4 | | E161 | | | | | | C105 | | |
| 5 | Y104 | E44 | | | | F32 | | S72 | | |
| 6 | | I46 | | | | | | C74 | | |
| 7 | | | K246 | | | | | R99 | | |
| 8 | | F279 | | | | | | G104 | | |
| 9 | D106 | | K247 | | | N33 | | G104 | | |
| 10 | Y107 | | K247 | | | | | C105 | | |
| 11 | W108 | | T70 | | | | | G106 | | |
| 12 | | | I113 | | | Y34 | | G102 | | |
| 13 | | | K246 | | | | | N103 | | |
| 14 | | | K247 | | | | | G104 | | |
| 15 | | | Q248 | | | S55 | R323 | | | |
| 16 | F109 | | S72 | | | R56 | Q325 | | | |
| 17 | | | V97 | | | | D362 | | | |
| 18 | | | R99 | | | S62 | D362 | | | |
| 19 | | | N103 | | | K68 | | | Q293 | |
| 20 | | | K246 | | | S69 | | | T176 | |
| 21 | L112 | | W101 | | | | | | G179 | |
| 22 | | | | | | | | | Q293 | |
| 23 | | | | | | G70 | | | G179 | |
| 24 | | | | | | | | | T180 | |
| 25 | | | | | | | | | K291 | |
| 26 | | | | | | | | | Q293 | |
| 27 | | | | | | N71 | | | E172 | |
| 28 | | | | | | | | | T180 | |
| 29 | | | | | | | | | K291 | |
| 30 | | | | | | S95 | | T70 | | |
| 31 | | | | | | | | A71 | | |
| 32 | | | | | | | | S72 | | |
| 33 | | | | | | R96 | | T70 | | |

**Table S12. Interacting residues between DENV2 N153Q and Fab C10 at 37ºC (Fab C10 binds to an epitope located near to 2-fold vertices)**

| No. | Fab C10 chain H | mol. B | mol. B' | mol. A | Fab C10 chain L | mol. B | mol. B' | mol. A |
|---|---|---|---|---|---|---|---|---|
| 1 | K59 | | T70 | | G31 | | C74 | |
| 2 | D103 | R2 | | | F32 | | S72 | |
| 3 | | I46 | | | | | C74 | |
| 4 | Y104 | H27 | | | | | R99 | |
| 5 | | | H244 | | | | G104 | |
| 6 | | | K246 | | | | C105 | |
| 7 | | F279 | | | N33 | | G104 | |
| 8 | D106 | | K247 | | | | C105 | |
| 9 | W108 | | T70 | | | | G106 | |
| 10 | | | I113 | | | K310 | | |
| 11 | | | K246 | | Y34 | | N103 | |
| 12 | | | K247 | | | | G104 | |
| 13 | | | Q248 | | D52 | K310 | | |
| 14 | F109 | | S72 | | | R323 | | |
| 15 | | | R99 | | T54 | V309 | | |
| 16 | | | N103 | | S55 | R323 | | |
| 17 | L112 | | W101 | | R56 | D362 | | |
| 18 | | | G102 | | S62 | D362 | | |
| 19 | | | G104 | | S69 | | | H52 |
| 20 | | | | | G70 | | | H52 |
| 21 | | | | | S95 | | T70 | |
| 22 | | | | | | | A71 | |
| 23 | | | | | | | S72 | |
| 24 | | | | | R96 | | T70 | |
| 25 | | | | | | | S81 | |
| 26 | | | | | | | L82 | |
| 27 | | | | | | | N83 | |
| 28 | | | | | | | | T226 |

**Table S13. Interacting residues between DENV2 N153Q and Fab C10 at 37ºC (Fab C10 binds to an epitope located near to 3-fold vertices)**

| No. | Fab C10 chain H | mol. C | mol. A' | mol. B | Fab C10 chain L | mol. C | mol. A' | mol. B |
|---|---|---|---|---|---|---|---|---|
| 1 | D102 | | K246 | | S26 | | R73 | |
| 2 | D103 | R2 | | | G31 | | C74 | |
| 3 | | E44 | | | F32 | | S72 | |
| 4 | | I46 | | | | | C74 | |
| 5 | Y104 | H27 | | | | | R99 | |
| 6 | | E44 | | | | | G104 | |
| 7 | | I46 | | | | | C105 | |
| 8 | | | K246 | | N33 | | G104 | |
| 9 | | F279 | | | | | C105 | |
| 10 | D106 | | K246 | | | | G106 | |
| 11 | | | K247 | | | K310 | | |
| 12 | W108 | | T70 | | Y34 | | N103 | |
| 13 | | | V97 | | | | G104 | |
| 14 | | | I113 | | D52 | K310 | | |
| 15 | | | K246 | | | R323 | | |
| 16 | | | K247 | | T54 | V309 | | |
| 17 | F109 | | S72 | | S55 | R323 | | |
| 18 | | | R99 | | R56 | Q325 | | |
| 19 | | | N103 | | | D362 | | |
| 20 | L112 | | W101 | | S62 | D362 | | |
| 21 | | | G102 | | S69 | | | H52 |
| 22 | | | G104 | | G70 | | | H52 |
| 23 | | | | | S95 | | T70 | |
| 24 | | | | | | | A71 | |
| 25 | | | | | | | S72 | |
| 26 | | | | | R96 | | T70 | |
| 27 | | | | | | | S81 | |
| 28 | | | | | | | L82 | |
| 29 | | | | | | | N83 | |
| 30 | | | | | | | | D225 |
| 31 | | | | | | | | T226 |

**Table S14. Number of interactions on the E protein – Fab interface.**

| DENV2-Fab complexes | Epitope location | No. of interactions | | | Interface area (Å$^2$) |
|---|---|---|---|---|---|
| | | Fab chain H | Fab chain L | E proteins | |
| DENV2 WT - Fab 2D22 at 37 °C | 5f | 38 / 9 | 10 / 9 | 48 / 18 (66) | 994 / 444 (1438) |
| | 2f | 45 / 0 | 12 / 0 | 57 / 0 (57) | 1082 / 77 (1159) |
| | 3f | 46 / 0 | 12 / 1 | 58 / 1 (59) | 1048 / 104 (1152) |
| DENV2 N153Q - Fab 2D22 at 37 °C | 5f | 43 / 6 | 13 / 7 | 56 / 13 (69) | 1028 / 399 (1427) |
| | 2f | 47 / 0 | 14 / 0 | 61 / 0 (61) | 1081 / 56 (1137) |
| | 3f | 46 / 0 | 10 / 0 | 56 / 0 (56) | 1030 / 54 (1084) |
| DENV2 WT - Fab C10 at 37 °C | 5f | NA | NA | NA | NA |
| | 2f | 21 / 0 | 22 / 2 | 43 / 2 (45) | 1067 / 116 (1183) |
| | 3f | 19 / 0 | 29 / 3 | 48 / 3 (51) | 1151 / 170 (1321) |
| DENV2 N153Q - Fab C10 at 37 °C | 5f | 21 / 0 | 20 / 13 | 41 / 13 (54) | 1106 / 173 (1279) |
| | 2f | 19 / 0 | 25 / 3 | 44 / 3 (47) | 1060 / 291 (1351) |
| | 3f | 22 / 0 | 27 / 4 | 49 / 4 (53) | 1079 / 169 (1248) |
| DENV2 NGC - Fab C10 (PDB ID: 7V3H) | 5f | 21 / 0 | 24 / 11 | 45 / 11 (56) | 1135 / 299 (1434) |
| | 2f | 23 / 0 | 21 / 3 | 44 / 3 (47) | 1108 / 134 (1242) |
| | 3f | 20 / 0 | 26 / 2 | 46 / 2 (48) | 1131 / 124 (1255) |
| DENV2 NGC - scFv C10 (PDB ID: 7CTH) | 5f | 16 / 0 | 13 / 21 | 29 / 21 (50) | 1036 / 292 (1328) |
| | 2f | NA | NA | NA | NA |
| | 3f | 15 / 0 | 23 / 1 | 38 / 1 (39) | 1066 / 113 (1179) |

The number of interactions and interface areas for interactions between Fab molecules and the E protein dimer or the adjacent E protein are separated by a forward slash (“/”). The total number of interactions and interface areas are shown in parentheses.
